# Development of lipidoid nanoparticles for siRNA delivery to neural cells

**DOI:** 10.1101/2021.07.28.454207

**Authors:** Purva Khare, Kandarp M. Dave, Yashika S. Kamte, Muthiah A. Manoharan, Lauren A. O’Donnell, Devika S Manickam

## Abstract

Lipidoid nanoparticles (LNPs) are the delivery platform in Onpattro, the first FDA-approved siRNA drug. LNPs are also the carriers in the Pfizer-BioNTech and Moderna COVID-19 mRNA vaccines. While these applications have demonstrated that LNPs effectively deliver nucleic acids to hepatic and muscle cells, it is unclear if LNPs could be used for delivery of siRNA to neural cells, which are notoriously challenging delivery targets. Therefore, the purpose of this study was to determine if LNPs could efficiently deliver siRNA to neurons. Because of their potential utility in either applications in the central nervous system and the peripheral nervous system, we used both cortical neurons and sensory neurons. We prepared siRNA-LNPs using C12-200, a benchmark ionizable cationic lipidoid along with helper lipids. We demonstrated using dynamic light scattering that the inclusion of both siRNA and PEG-lipid provided a stabilizing effect to the LNP particle diameters and polydispersity indices by minimizing aggregation. We found that siRNA-LNPs were safely tolerated by primary dorsal root ganglion neurons. Flow cytometry analysis revealed that Cy5 siRNA delivered via LNPs into rat primary cortical neurons showed uptake levels similar to Lipofectamine RNAiMAX—the gold standard commercial transfection agent. However, LNPs demonstrated a superior safety profile whereas the Lipofectamine-mediated uptake was concomitant with significant toxicity. Fluorescence microscopy demonstrated a time-dependent increase in the uptake of LNP-delivered Cy5 siRNA in a human cortical neuron cell line. Overall, our results suggest that LNPs are a viable platform that can be optimized for delivery of therapeutic siRNAs to neural cells.

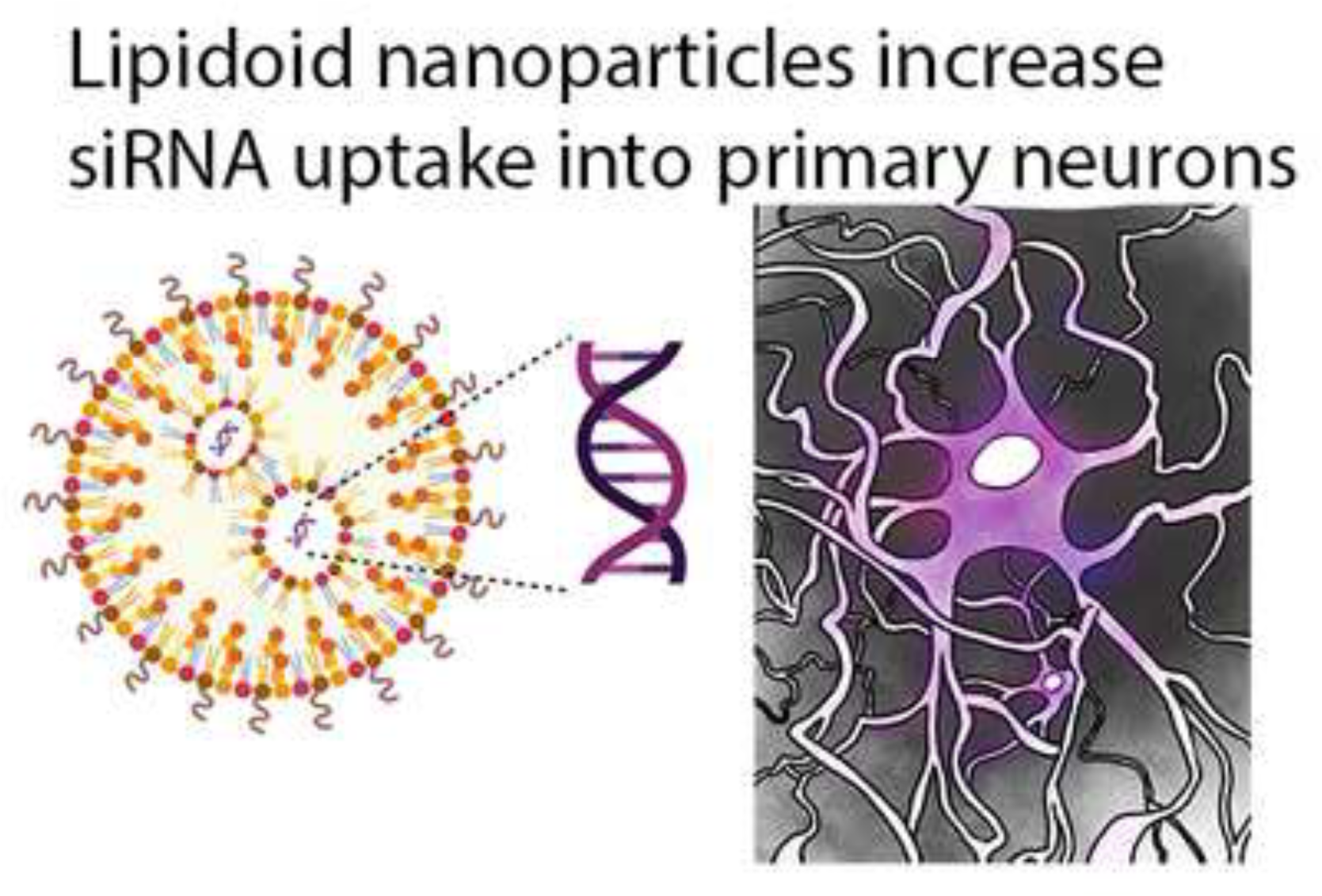
Graphical Abstract

## Introduction

Delivery of small interfering RNA (siRNA) is a promising strategy to treat pathologies as it allows genetic manipulation with a greater degree of target specificity with fewer off-target effects (1, 2). Lipidoid nanoparticles (LNPs) are the carriers in Onpattro, the first LNP-based RNAi drug to gain FDA approval for the treatment of polyneuropathies caused by a rare and life-threatening disorder, hereditary transthyretin-mediated amyloidosis (3). The clinical utility of LNPs is further reiterated by the recent full and emergency FDA approvals of the Pfizer-BioNTech (Comirnaty) and Moderna COVID-19 mRNA vaccines, respectively (4, 5). Based on the clinical success of LNPs in delivering siRNAs to hepatic and non-hepatic targets (7–9), we explored if LNPs could effectively deliver siRNA to neural cells. Neural cells are targets for drug delivery in multiple CNS disorders such as Alzheimer’s disease, Parkinson’s disease, ischemic stroke, peripheral nerve injuries, neuropathic and inflammatory pain (9–13). The key challenge associated with the delivery of siRNA to neural cells are its poor accumulation and short duration of action inside cells.

The potent delivery characteristics of LNPs may allow them to be optimized for neural cell delivery. LNPs are primarily composed of an ionizable cationic lipidoid along with helper lipids like distearylphosphatidylcholine (DSPC), polyethylene glycol-dimyristoyl glycerol (PEG-DMG) and cholesterol (13, 14). Ionizable cationic lipidoids allow higher encapsulation efficiencies and greater intracellular release via effective endosomal escape (16). Ionizable cationic lipidoids with pKa values between 6-7 acquire a positive charge at acidic pH and result in electrostatic interactions with the negatively charged siRNA molecules leading to the spontaneous formation of LNPs. At physiological pH of 7-7.4, these obtain a neutral charge and thereby largely eliminate the toxicity associated with cationic lipids such as lipofectamine (17). This key feature renders the LNPs a superior safety profile compared to cationic transfection agents. The acidic microenvironment of endosomes lend the LNPs a positive charge because of which they associate with negative anionic endosomal lipids leading to endosomal destabilization and improved siRNA release into the cytoplasm (17, 18).

Seminal studies by Langer, Anderson and colleagues developed high throughput lipidoid libraries consisting of thousands of ionizable cationic lipidoids and identified lead lipidoids based on their *in vitro* and *in vivo* delivery efficacies in rodents as well as non-human primates. Amongst those, C12-200 lipidoid demonstrated the highest knockdown and uptake *in vitro* (7, 19, 20). Therefore, we chose C12-200 lipidoid as the ionizable cationic lipidoid component of LNPs. LNP-based siRNA delivery harnesses several key advantages of LNP chemistry such as high transfection efficiency, low toxicity and immunogenicity, protection of the payload from physiological enzymes to increase their metabolic stability, ability for ligand conjugation and enhancing the overall pharmacokinetics of the delivered cargo (22–24). The novelty of our work lies in exploiting some of these cardinal features of LNPs to maximize siRNA uptake into neural cells, a notoriously challenging delivery target.

In this pilot study, we formulated siRNA-LNPs using C12-200 lipidoid and characterized their colloidal stability using dynamic light scattering. We characterized their cytocompatibility with a panel of breast cancer cell lines and a model of sensory neurons, primary dorsal root ganglion cultures. We studied the qualitative and quantitative uptake of siRNA delivered via LNPs into neurons using fluorescence microcopy and flow cytometry, respectively. Despite mediating similar levels of siRNA uptake, we found that LNPs are a safe carrier and show no signs of toxicity in the neuronal models compared to the commercially available cationic lipid, Lipofectamine. These findings encourage the further optimization of LNPs for knockdown of therapeutically-relevant neuronal targets.

### Experimental section

#### Materials

1,2-distearoyl-sn-glycero-3-phosphocholine (DSPC) (850365P) and 1,2-Dimyristoyl-rac–glycero-3-methoxypolyethylene glycol-2000 (PEG-DMG) (8801518) were obtained from Avanti Polar Lipids (Alabaster, AL). Cholesterol (8667) and Cy5-labeled siRNA were procured from Sigma-Aldrich (St. Louis, MO). 1,1‘-((2-(4-(2-((2-(bis (2 hydroxy dodecyl) amino) ethyl) (2hydroxydodecyl) amino) ethyl) piperazin1yl) ethyl) azanediyl) bis(dodecan-2-ol) (C12-200), an ionizable cationic lipidoid was a generous gift from Alnylam Pharmaceuticals (Cambridge, MA). Quant-iT Ribogreen RNA Assay Kit (R11490) was purchased from Life Technologies (Carlsbad, CA). Triton X-100 was procured from Acros organics (Morris Plains, NJ). Heat-inactivated fetal bovine serum (FBS) and phosphate buffered saline (PBS) were purchased from Hyclone Laboratories (Logan, UT). TrypLE express, RPMI + GlutaMAX-I (1X) and DMEM/F-12 were purchased from Life Technologies Corporation (Grand Island, NY). MCF-7, MDA-MB-231 and BT-549 cell lines were generously provided by Dr. Jane Cavanaugh (Duquesne University). Human cortical neuron cell line (HCN-2) (CRL-10742) and primary rat cortical neurons (RCN) (A10840-02) were purchased from ATCC (Manassas, VA) and Thermo Fisher Scientific (Frederick, MD) respectively. siGFP (AM4626), siGAPDH (Silencer Select GAPDH siRNA) (4390849), siTRPV1 (5’ GCGCAUCUUCUACUUCAACTT 3’, 3’ TTCGCGUAGAAGAUGAAGUUG 5’) [4390828 (Assay ID 554887)] and control, inverted siTRPV1 (5’ CAACUUCAUCUUCUACGCGTT 3’, 3’ TTGUUGAAGUAGAAGAUGCGC 5’) [4390828 (Assay ID 554886)] were procured from Thermo Fisher Scientific (Austin, TX). RNeasy Mini Kit (74104) and QIAshredder (79654) were procured from Qiagen (Qiagen, Germantown, MD). High-capacity cDNA Reverse Transcription Kit (4368813) was purchased from Thermo Fisher Scientific (ThermoFisher, Waltham, MA). Forward (AAGGATGGAACAACGGGCTAG) and reverse primers (TCCTGGTAGTGAAGATGTGGG) (299688565) for TRPV1 were procured from Integrated DNA Technologies (Coralville, Iowa). All reagents were used as received unless stated otherwise.

#### Preparation of siRNA-loaded LNPs (siRNA-LNPs)

LNPs were prepared using previously reported methods (8). Briefly, the ionizable cationic lipidoid C12-200, DSPC, cholesterol and PEG-DMG were dissolved in ethanol at a molar ratio of 50/10/38.5/1.5. The concentrations and volumes used for LNP preparation are shown in **Table 1**. A one mg/mL solution of siRNA was prepared in 10 mM citrate buffer, pH 4.0. The ‘slow mixing’ and ‘fast mixing’ protocols are defined based on the speed used during the addition of the lipid/ethanolic phase to the aqueous phase. For the ‘slow mixing’ method, the entire volume of the lipid phase was added to the aqueous phase instantaneously (in a single shot) followed by vortexing the mixture for 30 seconds while maintaining the Fisher benchtop vortexer knob at position ‘7’. For the ‘fast mixing’ protocol, the lipid phase was added dropwise to the aqueous phase under continuous vortexing for 30 seconds while maintaining the Fisher benchtop vortexer knob at position ‘7’. A precalculated volume of 1*X* PBS pH 7.4 was then added to the LNPs to adjust the final siRNA concentration to 400 nM. All the formulations were made at a final siRNA concentration of 400 nM and cationic ionizable lipidoid/siRNA w/w ratio was maintained at 5:1, unless stated otherwise.

**Table 1.**
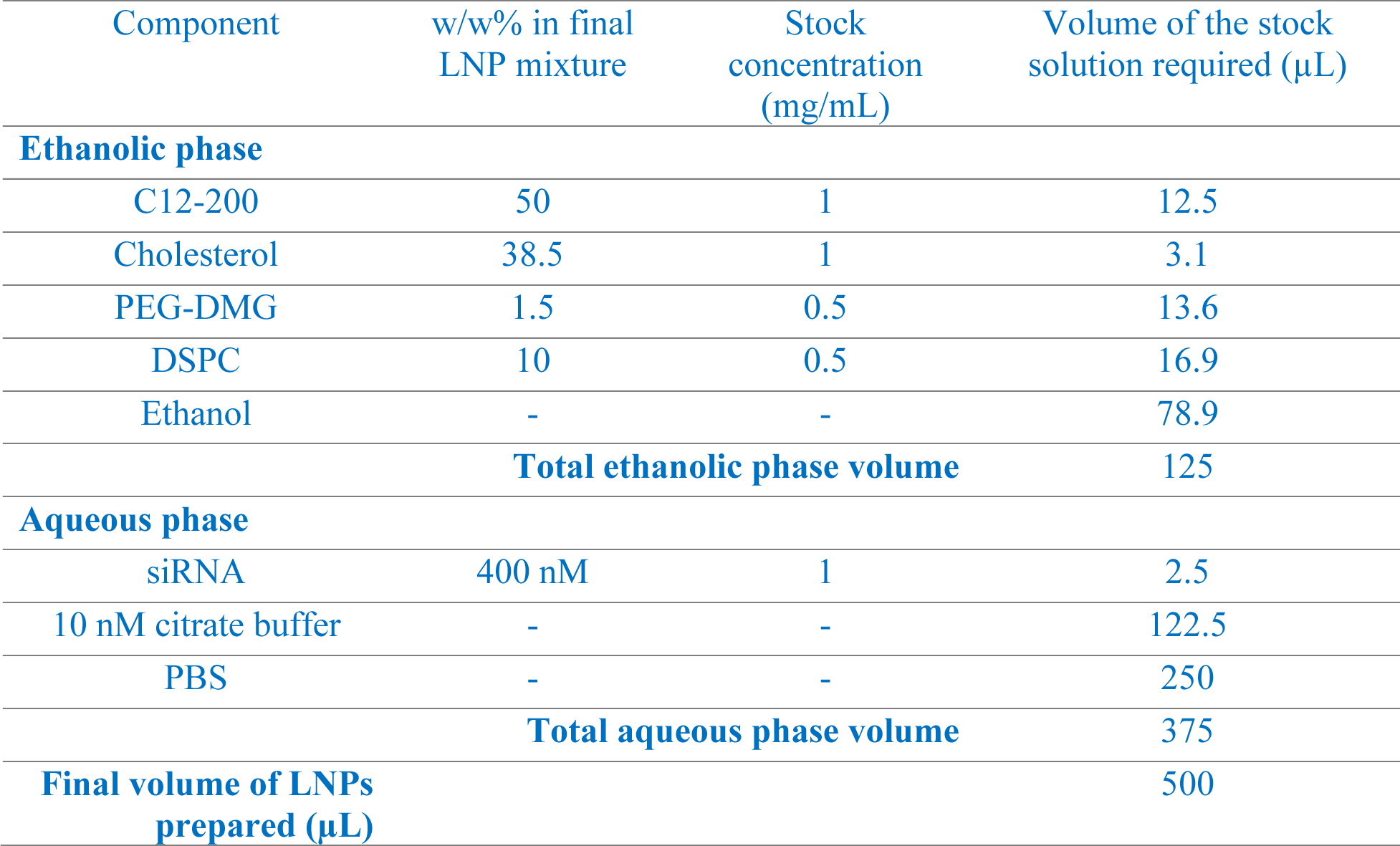
Representative formulation scheme for LNPs

| Component | w/w% in final LNP mixture | Stock concentration (mg/mL) | Volume of the stock solution required (μL) |
| --- | --- | --- | --- |
| **Ethanol phase** |  |  |  |
| C12-200 | 50 | 1 | 12.5 |
| Cholesterol | 38.5 | 1 | 3.1 |
| PEG-DMG | 1.5 | 0.5 | 13.6 |
| DSPC | 10 | 0.5 | 16.9 |
| Ethanol | - | - | 78.9 |
| **Total ethanol phase volume** |  |  | 125 |
| **Aqueous phase** |  |  |  |
| siRNA | 400 nM | 1 | 2.5 |
| 10 nM citrate buffer | - | - | 122.5 |
| PBS | - | - | 250 |
| **Total aqueous phase volume** |  |  | 375 |
| **Final volume of LNPs prepared (μL)** |  |  | 500 |

#### Agarose gel electrophoresis

Agarose gel electrophoresis was performed to qualitatively confirm the encapsulation of siRNA in LNPs. Briefly, a 2% agarose gel was prepared in 1*X* Tris-borate buffer composed of 108 g Tris and 55 g Boric acid dissolved in 900 ml distilled water with 40 ml of 0.5 M Na2EDTA and 0.05% Ethidium bromide. Samples pre-mixed with 1*X* RNA loading buffer were electrophoresed in an Owl EasyCast B2 Mini Gel Electrophoresis System (Thermo Fisher Scientific, St. Louis, MO) for 90 min. The gel was imaged under UV light using a Gel Doc system (ProteinSimple, San Jose, CA).

#### Determination of the extent of siRNA loading using the RiboGreen assay

The extent of siRNA loading in the prepared LNPs was measured using the Quant-iT Ribogreen RNA Assay Kit by following the manufacturer’s instructions. RiboGreen is a fluorescent dye, which, in its free form exhibits minimal fluorescence but when bound to a nucleic acid, it fluoresces with an exceptionally high intensity that is directly proportional to the amount of nucleic acid (25). Total siRNA content is defined as the amount of encapsulated and non-encapsulated/free siRNA in the LNPs. The difference between the total and free siRNA was used to calculate the amount of siRNA encapsulated within the LNPs. A fresh sample of two µg/mL dsRNA stock solution was prepared each time with 1*X* Tris-EDTA (TE) and the standard curve was generated at concentrations ranging from 0-1000 ng/mL. A free siRNA sample was prepared with 1*X* TE and LNPs were lysed with 2% Triton X-100 in 1*X* TE to measure the “total” siRNA content. The fluorescence intensities were measured on a SYNERGY HTX multi-mode reader (Winooski, VT) at an excitation wavelength of 485 nm and an emission wavelength of 528 nm. The % encapsulation efficiency was calculated using the **Eq. 1**.

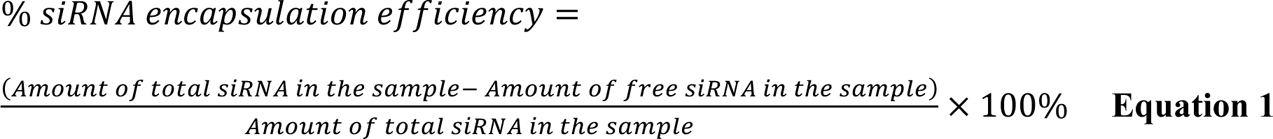

#### Dynamic Light Scattering

To determine the physical stability of LNPs, the average particle diameters, polydispersity indices and zeta potentials of the blank and siGFP-loaded LNPs were measured by dynamic light scattering (DLS) using a Malvern Zetasizer Pro (Malvern Panalytical Inc., Westborough, PA) over a period of seven days with intermittent storage at 2-8°C and upon storage at 37°C for 24 h. We also performed the colloidal stability analysis of LNPs 24 h post-preparation in 1*X* PBS and in samples supplemented with FBS (10%) to determine its serum stability. The prepared LNPs were diluted to a siRNA concentration of 40 nM using 10 mM 1*X* PBS pH 7.4 for particle size measurements and using deionized water for zeta potential measurements. All measurements were carried out in triplicate and the data are presented as average + standard deviation (SD) and are representative of at least five-six independent experiments (measurement errors <5%).

#### Cell culture

MCF-7 and BT-549 cells were maintained in RPMI + GlutaMAX (1*X*) supplemented with 10% FBS. MDA-MB-231 cells were maintained in DMEM/F-12 1:1 medium supplemented with 2.5 mM L-Glutamine + 15 mM HEPES Buffer + 110 mg/L Sodium Pyruvate + 10% FBS. The growth medium was changed every 48 hours. Primary rat dorsal root ganglion (DRG) cultures were isolated using previously reported methods (26). DRG cultures were maintained in DMEM/high glucose (4.5 g/L glucose) media supplemented with 2.5 mM L-Glutamine + 15 mM HEPES Buffer + 110 mg/L Sodium Pyruvate + 10 % FBS. Human cortical neuron cell line was maintained in DMEM/high Glucose (4.5 g/L glucose) medium supplemented with 10 % FBS. Primary rat cortical neurons were maintained in Neurobasal Plus medium supplemented with 200 mM GlutaMAX I and B-27 Plus supplement. All cultures were maintained in a humidified incubator at 37 °C and 5% CO_2_.

#### Cytocompatibility of LNPs

MCF-7, MDA-MB-231, BT-549 cells and primary rat DRG neurons were seeded in a clear, flat bottom Poly-D-Lysine coated 96-well plate (Azer Scientific, Morgantown, PA) for the cytocompatibility studies. MCF-7, MDA-MB-231, BT-549 cells were seeded at a density of 5,000 cells/well and DRG neurons were seeded at a density of 1,500 cells/well. Untreated cells were used as a control in all studies. Cells were transfected for 4 h with LNPs containing 50 nM of siRNA in complete growth medium in a volume of 50 μL/well. We also treated cells with increasing doses of siRNA-LNPs (10, 25, 50, 75 and 100 nM siGFP-loaded LNPs) to determine any possible effects of dose escalation on cytocompatibility. Post-transfection, the transfection mixture was replaced with 200 μL of fresh complete growth medium/well followed by a 24-hour incubation at 37°C and 5% CO_2_. The relative intracellular ATP levels were measured using the Cell Titer-Glo luminescence assay (ATP assay) 24 h-post transfection using previously described methods (27–29). The levels of ATP directly correlate with the cell numbers used here as a measure of cell viability. The ATP levels of the treatment groups were measured against those of the untreated or control cells. Briefly, 60 μL of the complete growth media and 60 μL of Cell Titer Glo 2.0 reagent were added to each well of the 96-well plate. The plate was incubated for 15 minutes in dark at room temperature in an incubator-shaker (Thermo Fisher Scientific, Waltham, MA). Following incubation, 60 μL of the mixture from each well was transferred to a flat-bottom, white opaque 96-well plate (Azer Scientific, Morgantown, PA). The luminescence was measured using SYNERGY HTX multi-mode reader (BioTek Instruments, Winooski, VT). Relative ATP levels (%) were calculated for the transfection groups by normalizing them to the luminescence of the untreated cells as shown in **Eq.2**.

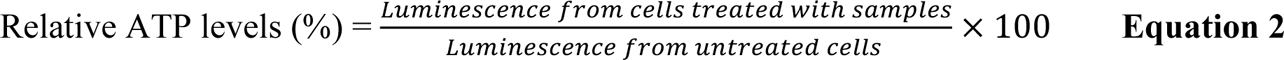

#### Cy5 siRNA uptake into primary rat cortical neurons using flow cytometry

Neurons were plated in a clear, flat-bottom Poly-D-Lysine coated 24-well plate (Genesee Scientific, San Diego, CA) at a density of 50,000 neurons/well in Complete Neurobasal Plus media and were allowed to acclimatize at 37°C and 5 % CO_2_ for 4-5 days. Cy5 siRNA containing LNPs were prepared as described earlier and diluted with the Complete Neurobasal Plus to a final Cy5 siRNA concentration of 50 nM. The neurons were incubated with the transfection mixture for 24 h. The transfection mixture contained either naked Cy5 siRNA or Cy5 siRNA-LNPs with PEG-DMG (+ PEG-DMG) or Cy5 siRNA-LNPs without PEG-DMG (-PEG-DMG) or Cy5 siRNA-Lipofectamine RNAiMAX complexes. Post-exposure, cells were gently removed from the wells and collected in microcentrifuge tubes. The cell suspension was centrifuged at 300 g for 5 minutes and the pellet was resuspended in 500 µL *1X* PBS. Cells were then analyzed using Attune NxT Acoustic Focusing Cytometer (Singapore) equipped with Attune NxT software. The fluorescence intensity of Cy5 siRNA was detected at an excitation wavelength of 638 nm and an emission wavelength of 720/30 nm. A total of 30,000 events were recorded for each sample. Histogram plots were obtained from the Attune NxT software. Cells were gated using forward vs. side scatter plots to exclude dead cells and cell debris. Untreated cells were used as negative-staining controls to set a gate for Cy5-negative and Cy5-positive populations. Data are presented as a percentage of Cy5-positive (Cy5 (+)) events.

#### Cy5 siRNA uptake by fluorescence microscopy in a human cortical neuron cell line

We used fluorescence microscopy to determine the Cy5 siRNA uptake into a neuronal cell line. Human cortical neurons (HCN-2) were seeded in a clear, flat-bottom, Poly-D-Lysine coated 48-well plate (Genesee Scientific, San Diego, CA) at a density of 7,000 cells/well. The cells were cultured in complete growth medium in a humidified incubator at 37°C and 5 % CO_2_ for 1-2 weeks while replenishing the growth medium every other day. After 4-5 days of culturing, the media was replaced with 175 μL of the transfection mixture. The transfection mixture consisted of either Cy5 siRNA-LNPs or Cy5 siRNA-Lipofectamine RNAiMAX complexes containing 50 nM or 100 nM of Cy5 siRNA or naked Cy5 siRNA diluted in complete growth media. Cells were incubated with the transfection mixture for two, four and 24 hours at 37°C and 5% CO_2_. After the respective time points, the cells were washed once using pre-warmed PBS before adding 0.5 mL of the complete, phenol red-free growth medium/well. Untreated cells and cells treated with Cy5 siRNA-Lipofectamine RNAiMAX complexes were used as negative and positive controls, respectively. Cells were imaged using an Olympus IX 73 epifluorescent inverted microscope (Olympus, Center Valley, PA) to detect Cy5 signals at excitation and emission wavelengths of 651 nm and 670 nm, respectively.

#### Transfection of siTRPV1-LNPs in primary rat DRG neurons

Primary rat DRG neurons were seeded in a clear, flat bottom Poly-D-Lysine coated 48-well plate (Azer Scientific, Morgantown, PA) at a density of 2,500 cells/well in triplicates. The cells were cultured in complete growth medium in a humidified incubator at 37°C and 5 % CO_2_ for 48 hours. Forty-eight hours post-seeding, the medium was replaced with 200 μL of the transfection mixture. Cells were transfected with either 50 nM naked siTRPV1, LNPs containing 50 nM of siTRPV1 or inverted siTRPV1 diluted in complete growth medium. Untreated cells and cells treated with RNAiMAX lipofectamine/siTRPV1 complexes were used as negative and positive controls, respectively. Cells were incubated with the transfection mixture for about eight-nine hours at 37°C and 5 % CO_2_. Post eight-nine hours of transfection, growth medium was added to bring up the volume 0.5 mL/well followed by a 24-hour incubation.

#### Quantitative reverse transcription PCR

Twenty-four hours-post transfection, the medium was removed, and cells were washed with 1*X* PBS and detached with TrypLE express. Detached cells were collected in microcentrifuge tubes and washed with 1*X* PBS. Total RNA extraction and purification from the cells was done using RNeasy Mini Kit and QIAshredder according to the manufacturer’s instructions. The isolated RNA was eluted in 30 µl of RNase-free water and stored at -80^°^C. The concentration of RNA was measured using NanoDrop ND-1000 spectrophotometer (Waltham, MA). Briefly, 1.5 µl of the sample was placed on the sample holder. The concentration (ng/µl) and the absorbance ratio of 260/280 were representative of the quantity and quality of the RNA, respectively. A 260/280 ratio of ∼2.0 was considered as “pure” for RNA. For mRNA detection, 2 µg of the RNA samples was reverse transcribed into cDNA in technical triplicates using a high-capacity cDNA Reverse Transcription Kit. All steps were done on ice to maintain integrity of the samples. Reverse transcription was done by a two-step protocol according to the manufacturer’s instructions. A DNA Thermal Cycler (Perkin Elmer Cetus) was used to incubate the samples. The resulting cDNA was diluted 10*X* with DEPC-treated water and stored at -20^°^C. Concentration of the cDNA was measured using NanoDrop ND-1000. One µg cDNA was used to perform RT-qPCR using TaqMan Gene Expression Master Mix and TaqMan Assays. An appropriate volume of the Gene Expression Master Mix and TaqMan primers were added to get a total volume of 20 µL. Reactions were performed in three technical replicates. The plate was covered with Microamp optical adhesive film and RT-qPCR was done using a Stratagene Mx3000P.

#### Statistical analysis

The data is expressed as mean ± standard deviation (SD), wherever applicable. Comparative statistical analyses were performed using either one-way, two-way ANOVA or One sample t and Wilcoxon tests using GraphPad Prism 9 (GraphPad Software, San Diego, CA). Bonferroni’s multiple comparisons test was performed for comparative analyses using one-way ANOVA, wherever applicable. Tukey’s and Šídák’s multiple comparisons tests for statistical comparisons were performed using two-way ANOVA, wherever applicable. Alpha was set at 0.05.

## Results

### Determination of the extent of siRNA loading using the RiboGreen assay

We used the Ribogreen assay as a quantitative measure to determine the extent of siRNA encapsulation within the LNPs ***(Figure 1)*** and used agarose gel electrophoresis as a qualitative tool to confirm the absence of ‘free’ or unentrapped siRNA **(*Supplementary Figure 1*)**. Triton X-100 was used to lyse the LNPs resulting in the extraction of the entrapped siRNA to measure the ‘total’ siRNA (Total siRNA = entrapped siRNA + free siRNA). Fluorescence readout from LNPs not treated with Triton X-100 is indicative of the amount of free siRNA. Two different mixing speeds used during LNP formulation, slow *vs*. fast, were compared for determining their effects on % siRNA encapsulation. The values increased from 74.4% at slow mixing to 83.8% at faster mixing speeds for LNPs prepared without the inclusion of PEG-DMG, whereas we did not observe a significant increase in siRNA loading for LNPs prepared with PEG-DMG at the different mixing speeds.

**Figure 1.**
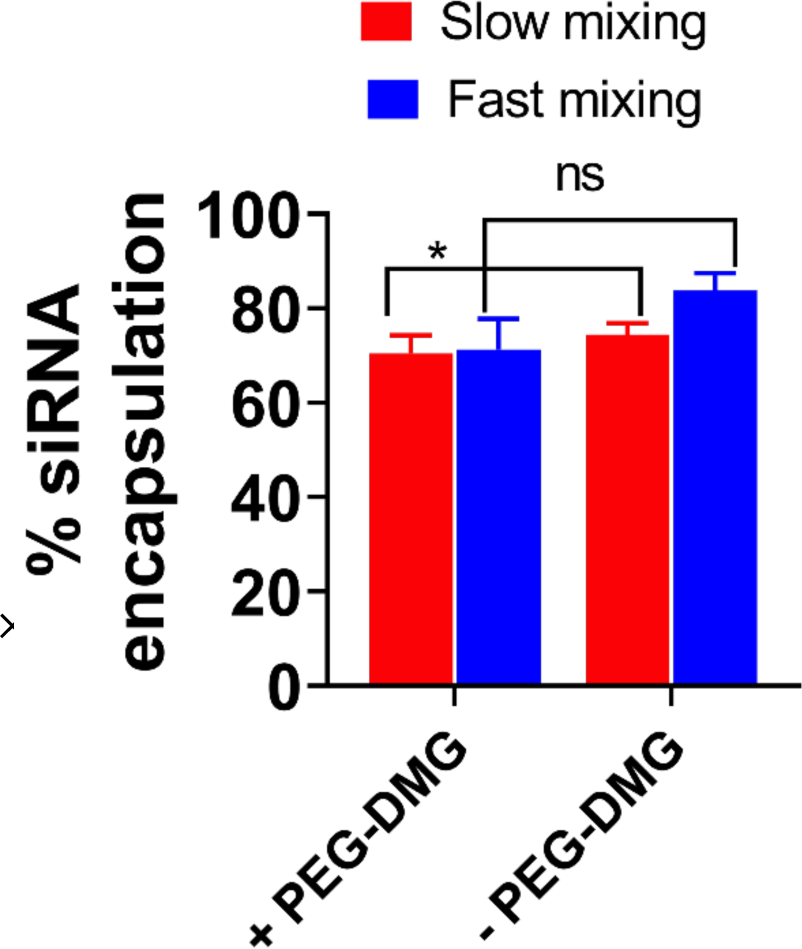
Extent of siRNA encapsulation in LNPs prepared at different mixing speeds determined by the Ribogreen assay. The extent of siRNA encapsulation in LNPs (+/- PEG-DMG) was calculated using the following equation (

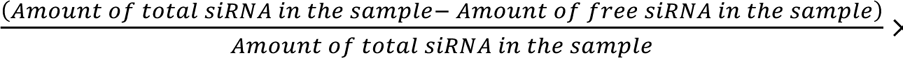

100%). SiRNA-LNPs were initially prepared in 10 mM citrate buffer and further diluted to a final siRNA concentration of 50 nM using 1*x* PBS. Data are presented as mean ± SD of n=3 samples. Statistical comparisons were made using two-way ANOVA. \**p* < 0.05 and ns: non-significant.

### Colloidal stability of siRNA-loaded LNPs

Physicochemical characteristics of nanocarriers such as particle diameter, polydispersity index (PDI) and surface charge (zeta potential) are influential determinants of their biological activity (30–32). Published studies have reported significant effects of physicochemical characteristics of nano-sized systems on the overall delivery efficacy (33–36). Particle characteristics of self-assembled systems such as LNPs can vary over time during storage potentially affecting their biological activity. We also anticipated that neural cells may show a slower rate of particle uptake compared to non-neuronal cells owing to limited endocytosis/other uptake pathways. We therefore measured the changes in particle diameters and zeta potentials of the LNPs over a period of seven days where the samples were refrigerated interim. Surface coating of nanoparticles using PEG-DMG has been reported to improve their pharmacokinetic profile by reducing the recognition by the mononuclear phagocyte system (37–39). We studied the effect of PEG-DMG in stabilizing LNPs by comparing LNPs prepared in the presence (+) and absence (-) of PEG-DMG. We compared the particle parameters of blank LNPs and siRNA-loaded LNPs to study the effect of siRNA encapsulation on the resulting particle characteristics.

Blank LNPs (+PEG-DMG) showed average particle diameters of about 250 nm which increased to about 350 nm on day seven. In contrast, the diameters of LNPs (-PEG-DMG) were about 900 nm post-preparation and increased to 4 µm after seven days ***(Figure 2a)***. A similar trend was observed for the PDI values of blank LNPs i.e., an initial PDI of 0.32 increased to 0.4 after seven days for LNPs (+PEG-DMG) and an initial PDI of 0.5 increased to 1.8 after seven days for LNPs (-PEG-DMG) ***(Figure 2b)***. We observed a few statistically significant changes in the particle diameters, PDI values and zeta potential of the prepared LNPs over the period of 7-days ***(Supplementary Table 1)***. The narrow polydispersity indices and smaller average particle diameters demonstrate the stabilizing effect of PEG-DMG in the LNPs. siGFP-loaded LNPs (with PEG-DMG) showed average particle diameters of about 120 nm which increased to about 200 nm on day seven ***(Figure 2a)***. Their initial PDI values of ca. 0.2 increased to about 0.3 after seven days ***(Figure 2b)***. From this observation we can infer that siRNA loading in the LNPs results in an additional stabilizing effect to the respective blank LNP counterparts resulting in narrow and consistent PDI values and particle diameters.

**Figure 2.**
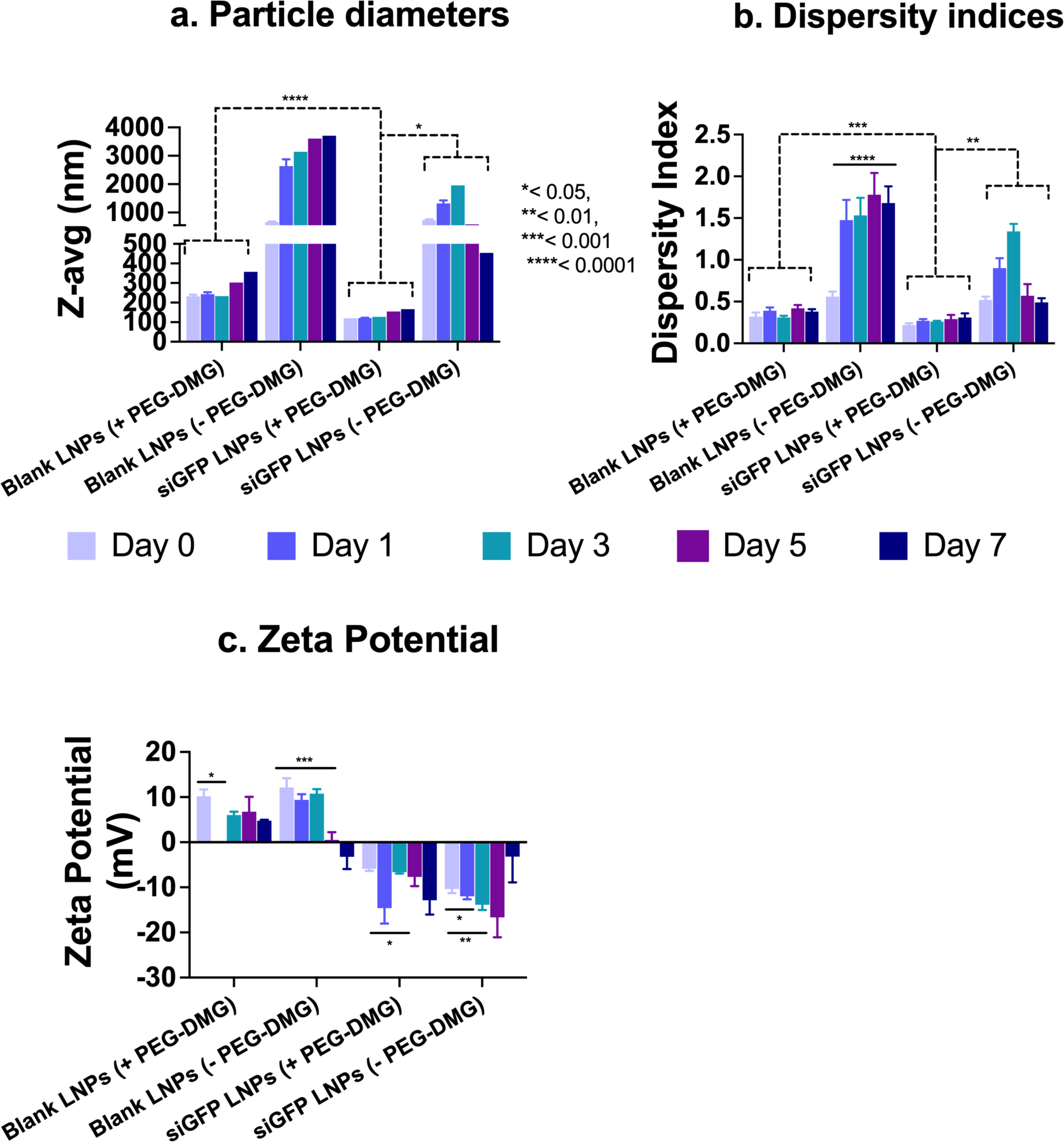
Colloidal stability of blank or siGFP-loaded LNPs measured using dynamic light scattering. SiGFP-LNPs were initially prepared in 10 mM citrate buffer and further diluted to a final siRNA concentration of 50 nM using 1*x* PBS (pH 7.4). The samples were stored at 2-8°C while not in use. Z-average particle diameters (a), dispersity indices (b) and zeta potentials (c) were measured on a Malvern Zetasizer Pro. Data are presented as mean ± SD of n=3 measurements. Statistical comparisons were made using one-way ANOVA or One sample *t* and Wilcoxon tests. \**p* < 0.05, \*\**p* < 0.01, \*\*\**p* < 0.001 and \*\*\*\**p* < 0.0001.

The cationic lipidoid, C12-200, in the LNPs resulted in a net positive charge to the blank LNPs. Blank LNPs containing PEG-DMG showed slightly lower zeta values (6-10 mV) compared to blank LNPs without PEG-DMG (9-12 mV) **(*Figure 2c*)** that is likely explained by the insertion of PEG chains in between the C12-200 lipidoids or due to formation of a PEG monolayer around C12-200 (40). As seen from ***Figure 2*c**, siGFP-loaded LNPs showed a negative zeta potential compared to their blank LNP counterparts. Although the changes were not statistically significant, the zeta potentials of the siGFP-loaded LNPs decreased over the seven-day period from -5.91 mV to -19.12 mV. Overall, the average particle diameters and polydispersity indices do not vary significantly over a period of seven days. However, zeta potential values of siGFP-LNPs decreased over a period of seven days. Despite the apparent numerical differences in the zeta potentials, a one-way ANOVA analysis revealed that the differences were not significant.

The colloidal stability of siGFP loaded- and blank LNPs was measured using dynamic light scattering at 0 (immediately after preparation) and 24 h post-preparation upon storage at 37°C **(*Figure 3*)**. We also studied the effect of PEG-DMG in stabilizing LNPs by preparing LNPs with (+) and without (-) PEG-DMG. Blank LNPs (+PEG-DMG) showed an average particle diameter of about 167.8 nm that shifted to about 159.8 nm 24 h-post storage at 37°C. In contrast, the diameters of LNPs (-PEG-DMG) were about 345.5 nm post-preparation and increased to 471.1 nm post 24 h storage at 37°C **(*Figure 3a*)**. A similar trend was observed for the PDI values of blank LNPs i.e., an initial PDI of 0.09 decreased to 0.06 for LNPs (+PEG-DMG) and an initial PDI of 0.3 increased to 0.4 for LNPs (-PEG-DMG) 24 h-post storage at 37°C **(*Figure 3b*)**. We found a similar trend for the particle diameters and dispersity indices of siGFP-loaded LNPs, both (+) and (-) PEG-DMG, 24 h-post storage at 37°C **(*Figure 3a and b*)**. siGFP-loaded LNPs (+PEG-DMG) showed an average particle diameter of about 163.2 nm that shifted to about 157.2 nm 24 h-post storage at 37°C. In contrast, the diameters of LNPs (-PEG-DMG) were about 442.2 nm post-preparation and decreased to 390.7 nm 24 h-post storage at 37°C **(*Figure 3a*)**. For the dispersity indices, an initial value of 0.09 decreased to 0.06 for siGFP-loaded LNPs (+PEG-DMG) and an initial value of 0.31 shifted to 0.32 for LNPs (-PEG-DMG) 24 h-post storage at 37°C **(*Figure 3b*)**. Nevertheless, the observed changes in particle diameters and dispersity indices of blank LNPs (+ PEG-DMG) and siGFP-loaded LNPs (+ and – PEG-DMG) 24 h-post storage at 37°C were statistically insignificant. However, we noted a significant increase in the particle diameters (*p < 0.05) and dispersity indices (****p < 0.0001) of blank LNPs (-PEG-DMG) likely due to the absence of PEG-DMG and the siGFP cargo—that are known to provide a stabilizing effect by inhibiting particle-particle aggregation and via complexation of the negatively charged siRNA with the positively-charged C12-200, respectively. Noteworthy, we noted a similar trend of the stabilizing effects of PEG-DMG and the siGFP cargo in ***Figure 2***.

**Figure 3.**
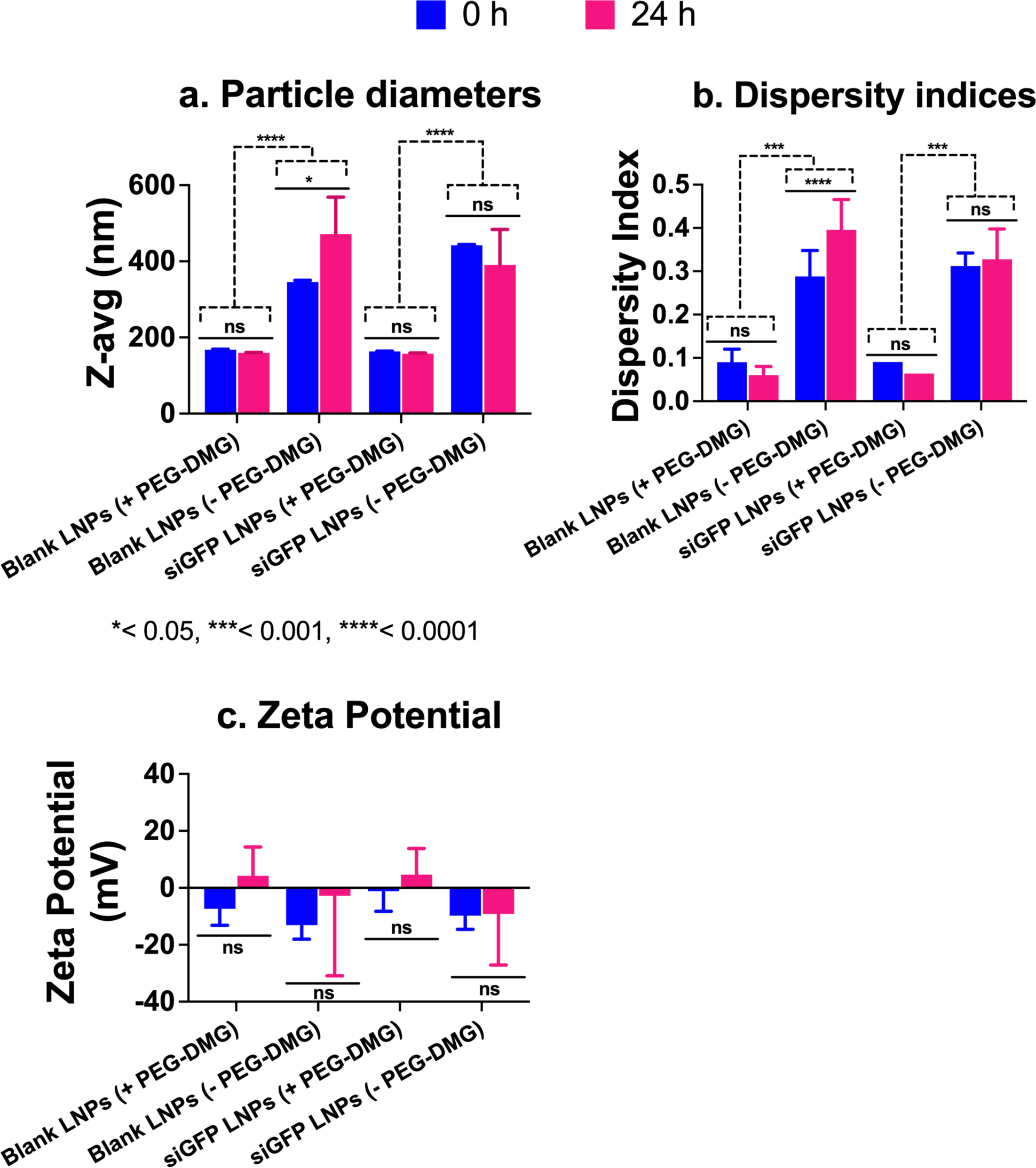
Colloidal stability of blank- or siGFP-loaded LNPs immediately after preparation (0 h) and upon storage at 37°C for 24 h measured using dynamic light scattering. SiGFP-LNPs were initially prepared in 10 mM citrate buffer and further diluted to a final siRNA concentration of 50 nM using 1x PBS at a final pH of 7.4 for particle size and PDI measurements and using de-ionized water for zeta potential measurements. The samples were stored at 37°C for 24 h. Z-average particle diameters (a), polydispersity indices (b) and zeta potentials (c) were measured on a Malvern Zetasizer Pro. Data are presented as mean ± SD of n=3 measurements. Tukey’s and Šídák’s multiple comparisons tests for statistical comparisons were performed using two-way ANOVA. *p < 0.05, ***p < 0.001, ****p < 0.0001 and ns: non-significant.

Blank LNPs containing PEG-DMG showed slightly negative zeta potentials (-5 to -7 mV) that shifted to slightly cationic values (3 to 4 mV) whereas the zeta potentials of blank LNPs without PEG-DMG were about -13 mV that increased to about -2 mV 24 h-post storage at 37°C **(*Figure 3c*)**. siGFP-loaded LNPs (+ PEG-DMG) showed a zeta potential of -1.2 mV that increased to about 4.6 mV whereas the LNPs (-PEG-DMG) counterparts showed a zeta potential of -9.8 mV that increased to about -9.1 mV 24 h-post storage at 37°C **(*Figure 3c*)**. Despite the apparent numerical differences, it should be noted that the measured zeta potentials reflect electrostatically-neutral samples and a two-way ANOVA analysis revealed that the differences were not significant.

The particle diameters of siGFP-LNPs (+PEG-DMG) containing 10% FBS was measured using dynamic light scattering to determine the effect of serum proteins on the colloidal stability of LNPs. We specifically chose siGFP-LNPs (+PEG-DMG) to study the effect of serum since the presence of siRNA and PEG-DMG resulted in the maximum stability of LNPs with respect to particle diameters and dispersity indices **(Figure 2*a* and *b*)**. We first measured the particle diameter of a control sample containing 10% FBS in PBS **(*Figure 4a*)** that showed an average particle diameter of 19.6 nm whereas siGFP-LNPs (+PEG-DMG) in PBS showed an average particle diameter of about 170 nm ***(Figure 4b)***. We observed distinct size distribution by intensity peaks for the blank 10% FBS as well as LNP samples ***(Figure 4a and b)***. We then measured the particle diameter of siGFP-LNPs (+PEG-DMG) that was supplemented with 10% FBS. ***Figure 4*c** has two distinct peaks denoting peaks corresponding to 10% FBS and the siGFP-LNPs (+PEG-DMG) at 20.2 nm and 80.5 nm respectively. The overlay size distribution plot demonstrated a “shift” in the siGFP-LNPs (+PEG-DMG) peak towards the left (lower particle diameters) when the LNPs were supplemented with 10% FBS as opposed to those in PBS (80.5 nm vs 170 nm) **(*Figure 4d)***. We speculate that the observed decrease in diameter is suggestive of the additional stabilizing effects of serum (10% FBS) on the LNP diameters. This, in fact, may be beneficial/favorable for LNP uptake ***(Figure 7 and Figure 8)*** as neurons may preferentially internalize smaller compared to larger LNPs.

**Figure 4.**
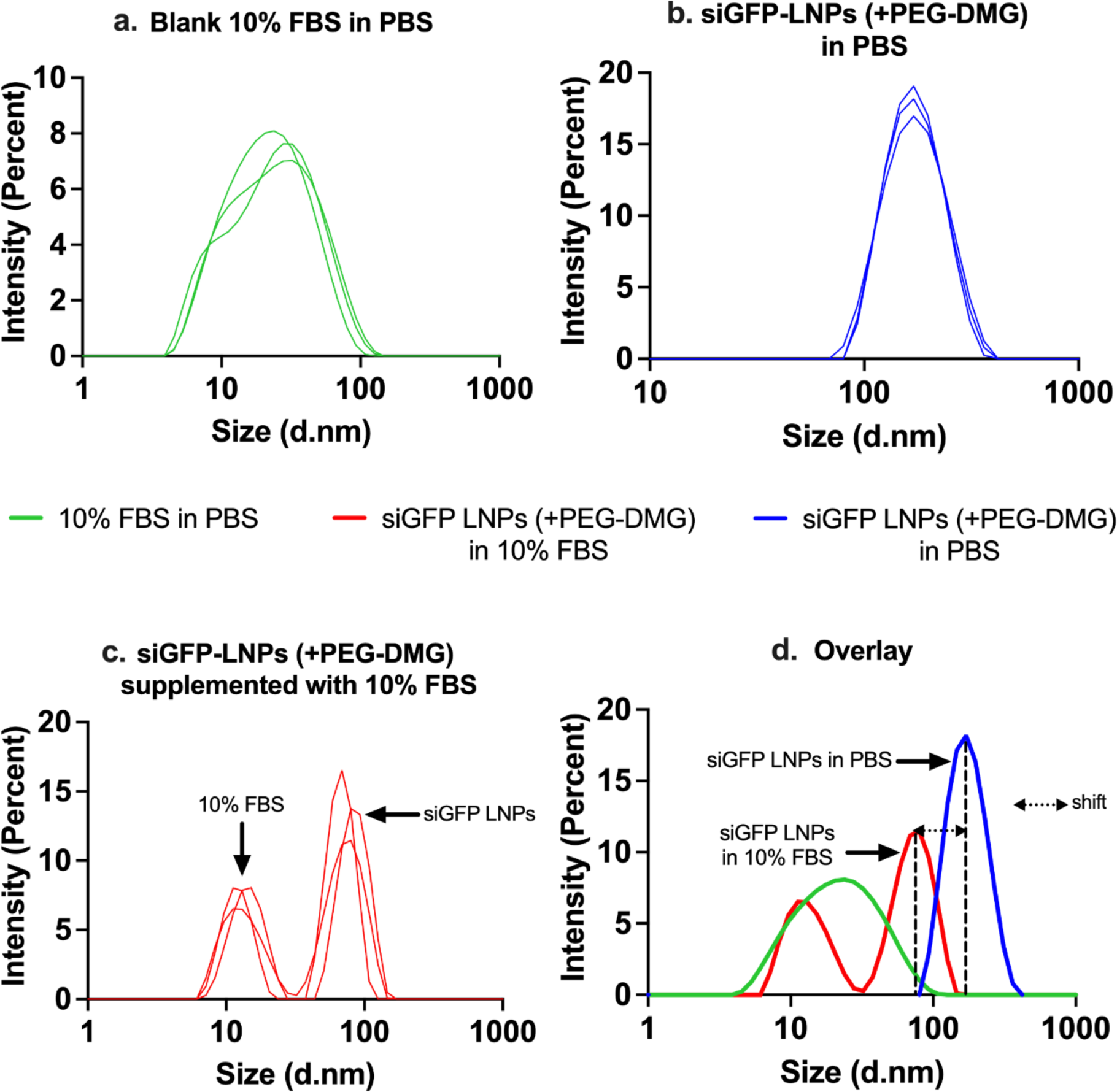
Intensity size distribution plots demonstrating the colloidal stability of siGFP-loaded LNPs (+PEG-DMG) supplemented with 10% FBS. (a) A blank sample consisting of 10% FBS in 1x PBS was used to understand the serum protein-mediated scattering. (b) SiGFP-LNPs (+PEG-DMG) were initially prepared in 10 mM citrate buffer and further diluted to a final siRNA concentration of 50 nM using 1x PBS. LNPs supplemented with 10% FBS were measured to determine their serum stability. (d) An overlay depicts the pattern of the size distribution obtained for all the LNP samples.

The correlation function is a statistical analysis tool for measuring the non-randomness in a data set that is depicted here as a correlogram. A correlogram is a plot of the correlation coefficient G (τ) vs time (µs) (**Figure 5**). The Y-intercept of the correlogram is indicative of the signal to noise ratio and presence/absence of multiple scattering. A Y-intercept of ∼0.9-1.15 indicates a good signal in the absence of multiple scattering. Samples with multiple scattering show a Y-intercept of ∼0.6-0.8 and samples with number fluctuations show a Y-intercept of ∼1.15-1.4 (41). As observed from **Table 2**, the Y-intercept values for all the LNP groups over a period of seven days ranged from 0.8-1.1 that is indicative of a good signal-to-noise ratio in the absence of multiple scattering and number fluctuations.

**Figure 5.**
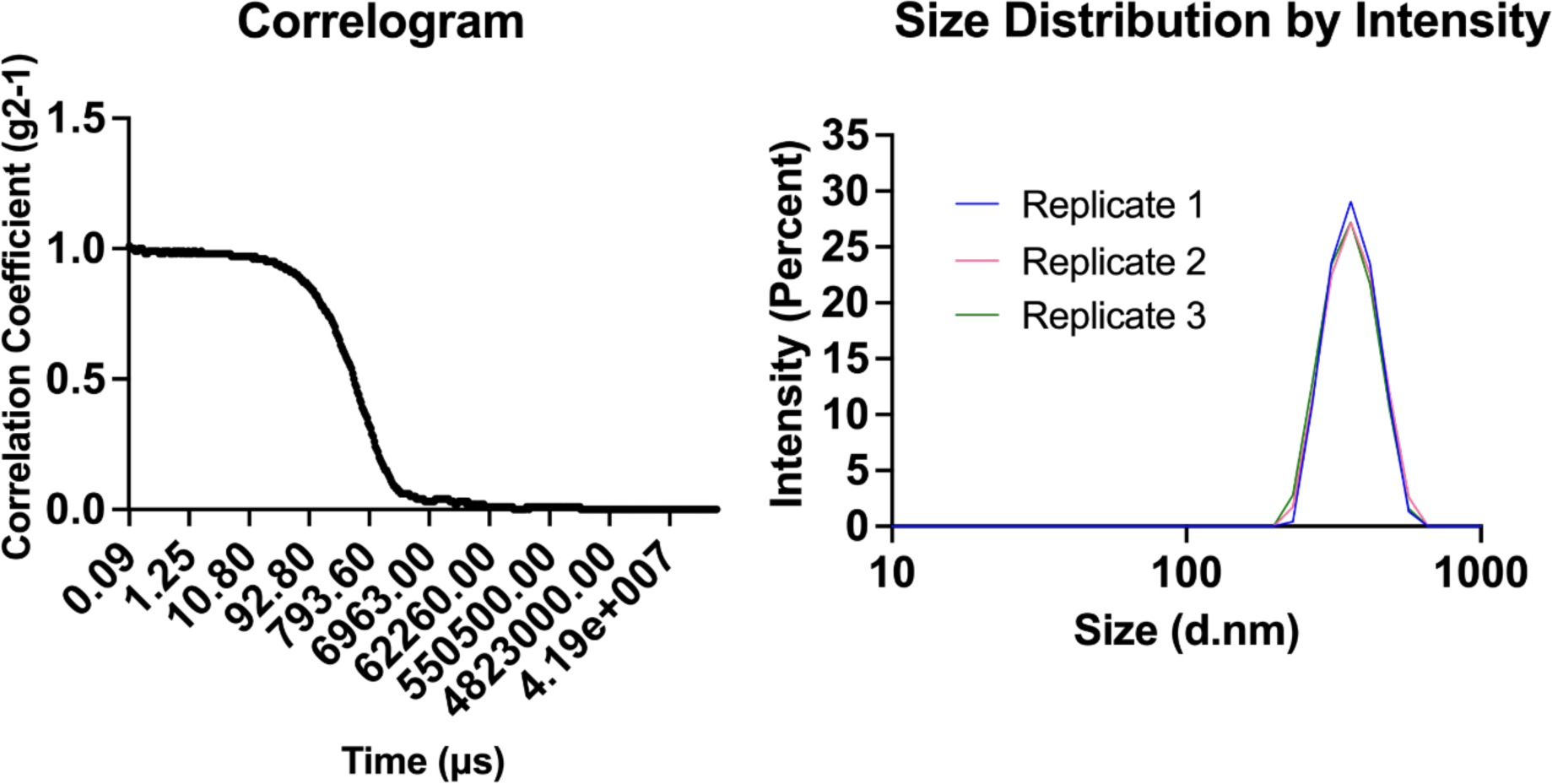
Representative correlogram and intensity size distribution plots of siGFP-loaded LNPs (+PEG-DMG). siGFP-LNPs were initially prepared in 10 mM citrate buffer and further diluted to a final siRNA concentration of 50 nM using 1*x* PBS at a final pH of 7.4 prior to particle size measurements on a Malvern Zetasizer Pro.

**Figure 6.**
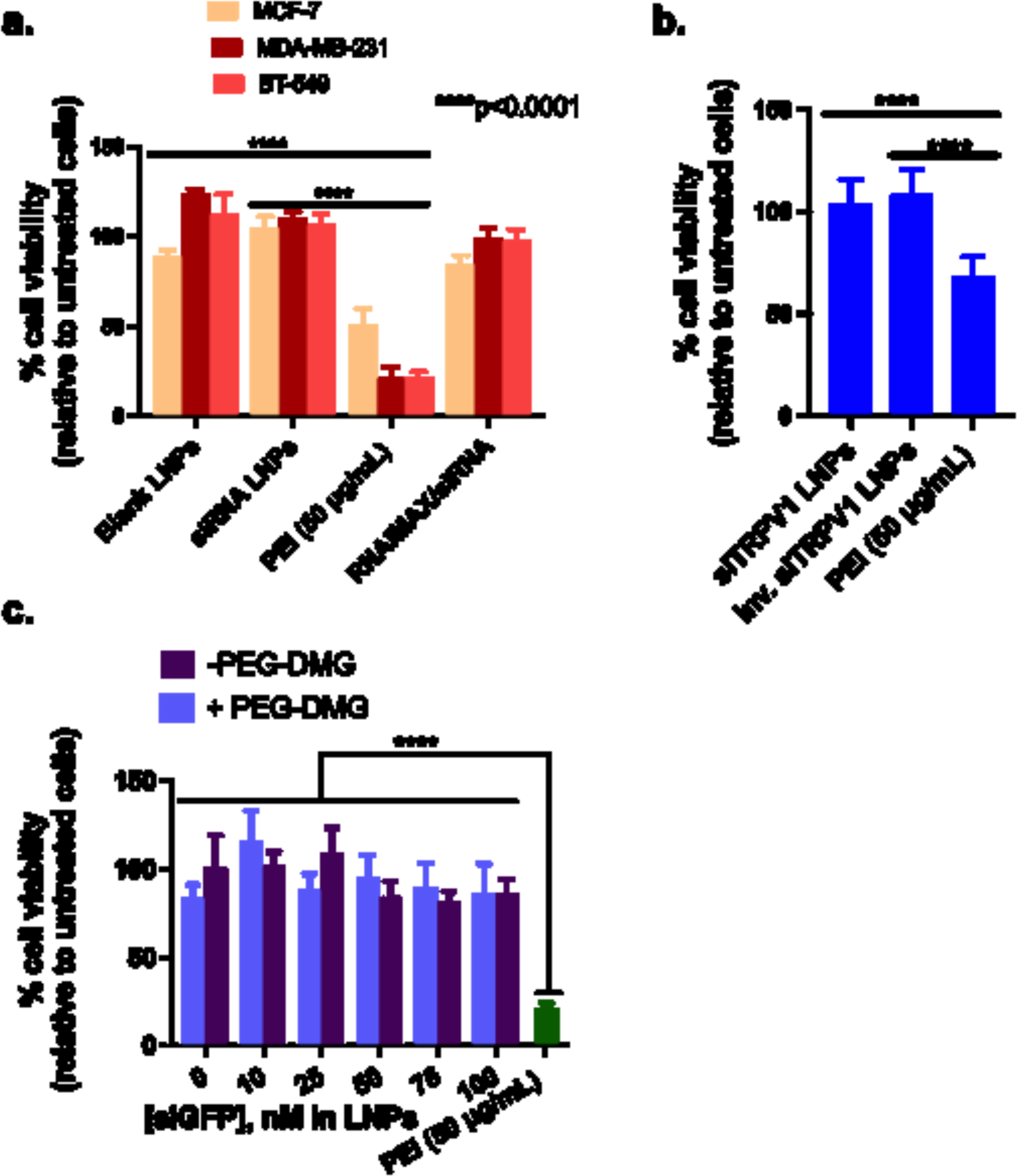
C12-200 LNPs are cytocompatible with TRPV1-expressing breast cancer cell line panel (a), rat primary DRG neurons (b) and MDAMB- 231 cells (c). Cells were incubated with LNPs containing 50 nM siGAPDH (a), 50 nM siTRPV1 (b) and the indicated concentrations of siGFP-LNPs (c) for four hours. Untreated cells and cells treated with PEI (50 μg/mL) were used as negative and positive controls, respectively. Cell viability was determined 24 h-post transfection using a CellGlo luminescence viability assay and the data were normalized to untreated cells. Data represents mean + SD (n=6).

**Figure 7.**
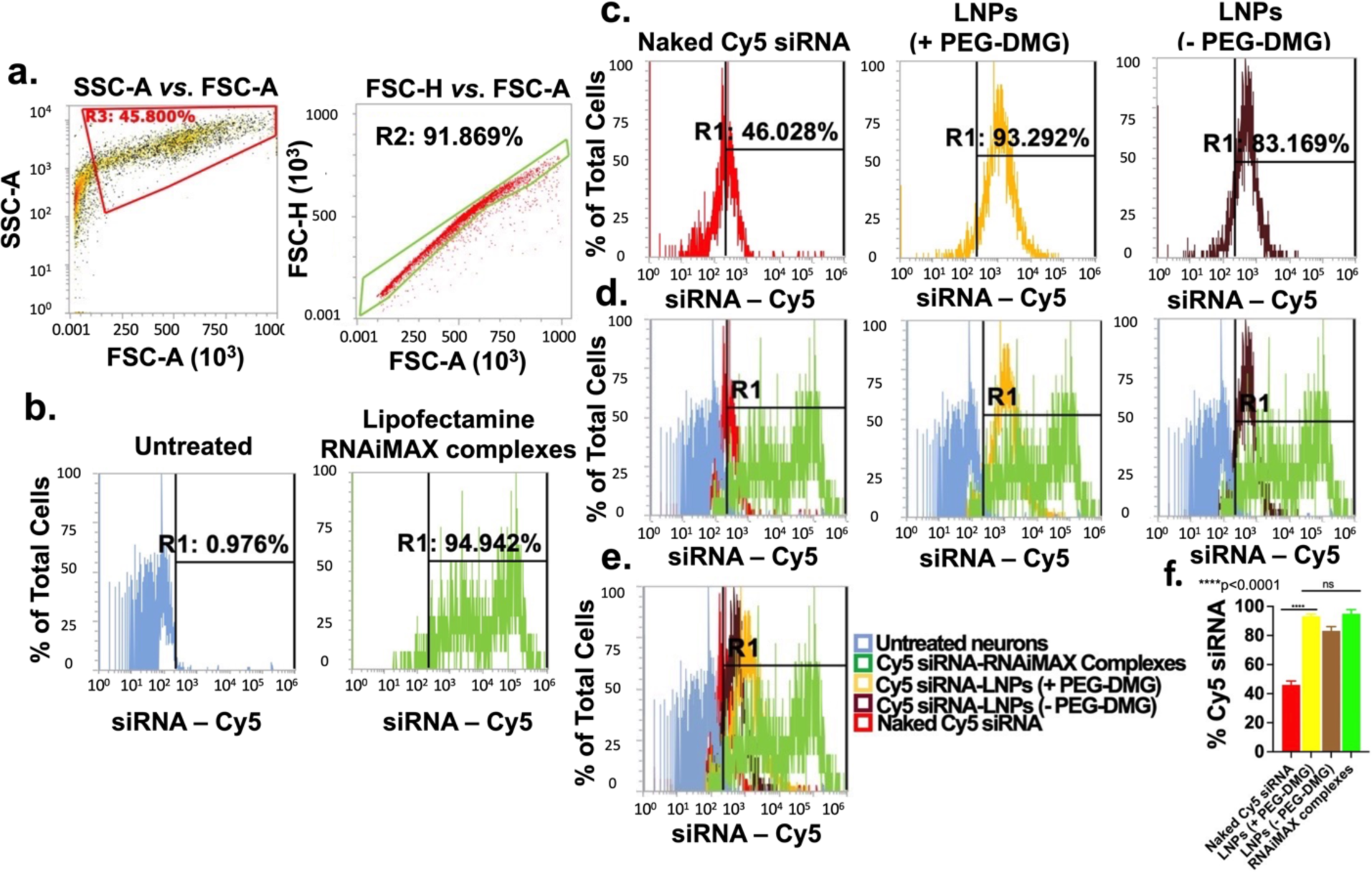
Uptake of LNPs into rat primary cortical neurons determined using flow cytometry analysis. Cells were transfected for 24 h with the indicated samples in a 24-well plate. Cy5 siRNA-Lipofectamine RNAiMAX complexes and untreated cells were used as positive and negative controls, respectively. Data are presented as percentages of positive cells for Cy5 (Cy5 (+)). SSC-A vs. FSC-A and FSC-H vs. FSC-A plots of untreated cells demonstrating monodispersed cells. (a), Untreated and Cy5 siRNA/Lipofectamine RNAiMAX-treated cells (b), Cells treated with the indicated samples (c), Overlay histograms comparing Cy5 (+) cells in each treatment group in comparison to the controls (d), Overlay histograms (e) and % Cy5 siRNA uptake by the neurons for the respective groups (f). The histograms are representative of quadruplicate samples. Statistical comparisons were made using one-way ANOVA. ****p < 0.0001.

**Figure 8.**
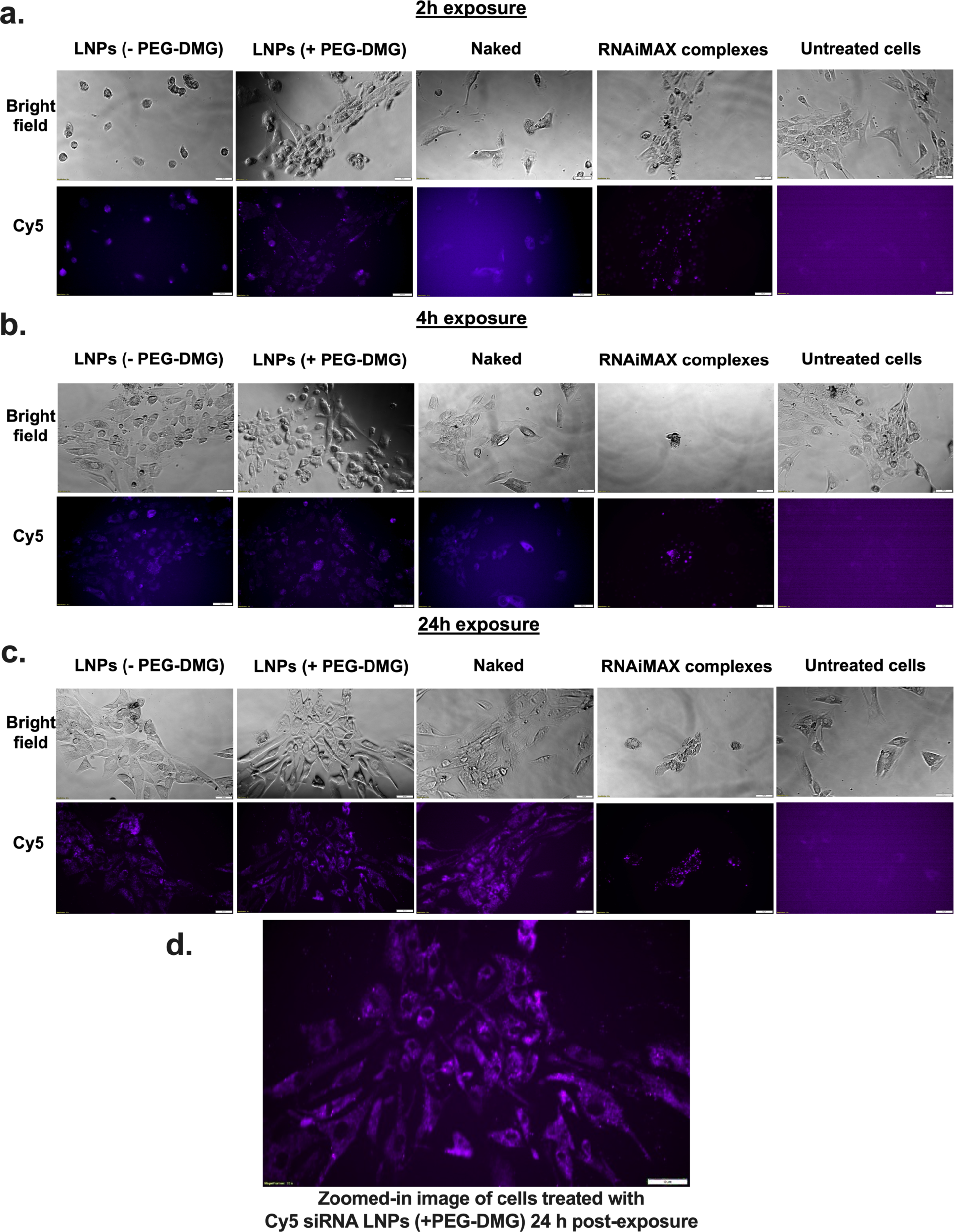
Cellular uptake of Cy5 siRNA determined using fluorescent microscopy. HCN-2 cells (human cortical neuron cell line) were incubated for 2 hours (a) 4 hours (b) and 24 hours (c) with the indicated samples containing 50 nM Cy5 siRNA. Panel (d) represents the zoomed-in image depicting HCN-2 cells incubated with LNPs with PEG-DMG. Scale bar = 50 µm. Cells were imaged using an Olympus IX 73 epifluorescent inverted microscope to detect Cy5 signals at excitation and emission wavelengths of 651 nm and 670 nm, respectively. Images are representative of n=3 independent wells.

**Table 2.**
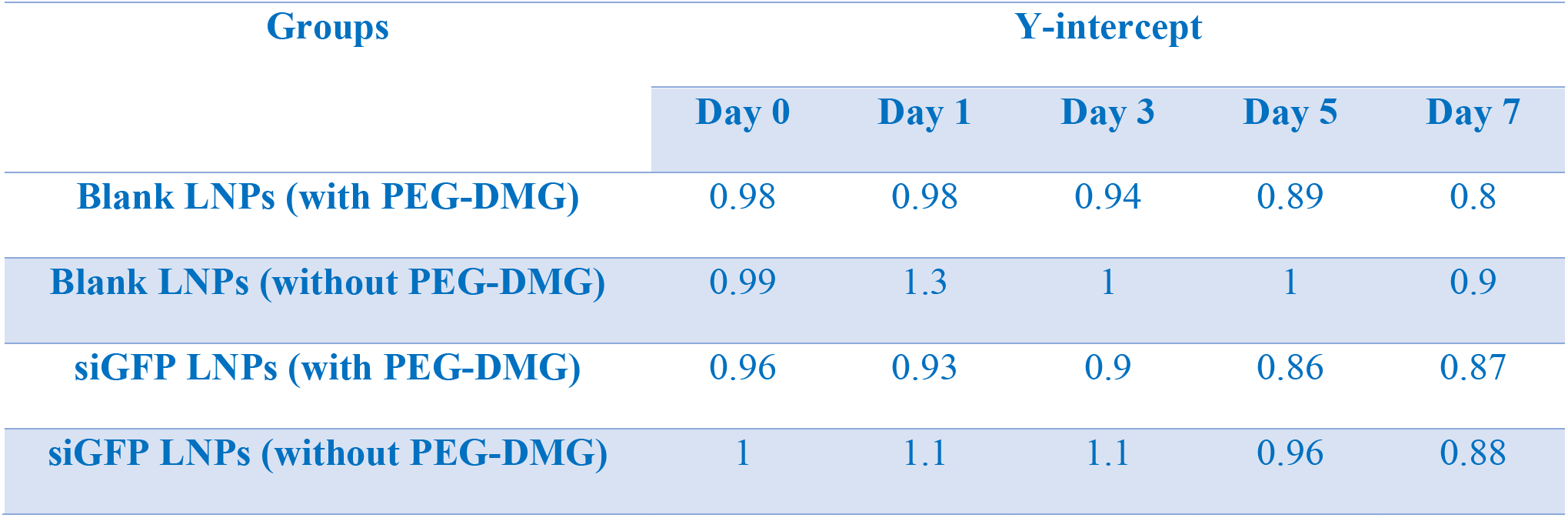
Comparison of the Y-intercept in the autocorrelograms from the colloidal stability data (Figure 2) at different time points.

| Groups | Y-intercept |  |  |  |  |
| --- | --- | --- | --- | --- | --- |
|  | Day 0 | Day 1 | Day 3 | Day 5 | Day 7 |
| Blank LNPs (with PEG-DMG) | 0.98 | 0.98 | 0.94 | 0.89 | 0.8 |
| Blank LNPs (without PEG-DMG) | 0.99 | 1.3 | 1 | 1 | 0.9 |
| siGFP LNPs (with PEG-DMG) | 0.96 | 0.93 | 0.9 | 0.86 | 0.87 |
| siGFP LNPs (without PEG-DMG) | 1 | 1.1 | 1.1 | 0.96 | 0.88 |

### Cytocompatibility of C12-200-LNPs

We studied the cytocompatibility of the prepared C12-200/siGAPDH LNPs with MCF-7, MDA-MB-231 and BT-549 cells as these cell models are known to overexpress TRPV1, a neuronal target for siRNA delivery ***(Figure 6a)*** (42). We also studied the compatibility of C12- 200/siTRPV1 LNPs with primary rat dorsal root ganglion (DRG) neurons ***(Figure 6b)***. We specifically chose primary rat DRG neurons due to their high expression of TRPV1 (41, 42). TRPV1 is reported to be expressed in 60% of the peripheral nociceptors present in the DRG (45) and is a potential neuronal target for gene knockdown. Cell Titer Glo assay (referred henceforth as the ATP assay) was used the study cytocompatibility of LNPs. The ATP assay is a simple and straightforward technique wherein the readout in relative luminescence units (RLU) can be directly correlated with the ATP levels in the cells. RLU of the untreated cells were normalized to 100% cell viability. Untreated cells and cells treated with a synthetic polycation, polyethyleneimine (PEI) served as the negative and positive controls, respectively. As we expected, MCF-7, MDA-MB- 231, BT-549 cells and primary DRG neurons treated with PEI (50 μg/mL), a synthetic polycation known to induce apoptosis showed a 50-60% reduction in cell viability (46). However, MCF-7, MDA-MB-231 and BT-549 incubated with C12-200/siGAPDH LNPs showed very little evidence of cell death, with cell viabilities >95% 24 h-post transfection ***(Figure 6a)***. Similarly, siTRPV1- loaded C12-200 LNPs were well tolerated (cell viabilities of ca. 100%) by primary DRG neurons ***(Figure 6b)***. We tested the effect of siRNA dose escalation by incubating MDA-MB-231 cells with increasing concentrations of siRNA-LNPs (10, 25, 50, 75 and 100 nM siGFP-loaded LNPs). We noticed cell viabilities >85% 24 h-post transfection **(*Figure 6c)*** suggesting that higher siRNA and concomitant lipid doses were well-tolerated by the cells. There was a significant difference (p<0.0001) in the % cell viabilities of cells treated with siRNA-LNPs compared to those treated with the positive control, PEI (20% cell viability).

### Flow cytometry analysis of Cy5 siRNA uptake into primary rat cortical neurons

After confirming that LNPs had little-to-no effect on cell viability, we next wanted to determine if LNPs were taken up by neurons. Because of their commercial availability as well as their non-mitotic nature similar to that of the primary rat DRG neurons, primary rat cortical neurons were used to quantify the uptake levels of Cy5 siRNA using flow cytometry. Lipofectamine RNAiMAX, a cationic lipid, was used as a positive control. Flow cytometry is a quantitative tool for single-cell analysis to measure the light scattered and the fluorescence emitted form a single cell (47). The dead cells/debris were likely removed at the centrifugation step during the sample processing and as a result, we did not observe any dead cells or debris ***(Figure 7a, left)*** that are typically found at the bottom left corner of the SSC-A vs. FSC-A plot (48). We then analyzed whether the cells were present in a monodisperse cell suspension to allow for single cell analysis. The forward scatter distributions for height *vs*. area (FSC-A v/s FSC-H) showed a linear profile for about 92% of the recorded events which indicated a single-cell suspension ***(Figure 7a, right)***. We analyzed untreated cells and gated out the auto fluorescence ***(Figure 7b, left)***. Thus, a shift of signal to the right of the histogram gate (R1 region) was considered to signify that the cell was Cy5-positive (+). The proportion of Cy5 (+) cells directly correspond to the % uptake of Cy5 siRNA in the neurons as represented by the individual histogram plots for the different groups ***(Figure 7c)***.

We then analyzed the % uptake of Cy5 siRNA encapsulated in LNPs prepared with (+) and without (-) the inclusion of PEG-DMG. Around 92-94% ***(Figure 7c (yellow))*** and 81-83% ***(Figure 7c (brown))*** cells were Cy5 (+) when transfected with Cy5 siRNA-LNPs prepared with and without the inclusion of PEG-DMG, respectively. ***Figure 7*d** shows the relative Cy5+ cells in each of the indicated groups in reference to the untreated (blue) and positive control-treated (green) groups. The rightward shift in intensities of Cy5 (+) cells for all the groups in comparison with the controls are demonstrated in the overlay histogram plot in ***Figure 7*e**. As expected, the % uptake of Cy5 siRNA was least in cells treated with free Cy5 siRNA and greatest in cells treated with LNPs (+PEG-DEMG) and Cy5 siRNA/Lipofectamine RNAiMAX complexes ***(Figure 7e and f)***. Furthermore, as seen in ***Figure 7*e**, we saw a greater uptake of LNPs with PEG-DMG as compared to those without PEG-DMG. Particle sizes of LNPs play one of the major roles in determining uptake levels and therefore, lower diameter particles may allow efficient cellular internalization. Furthermore, a significant difference in the uptake of Cy5 siRNA was observed when transfected with naked Cy5 siRNA and Cy5 siRNA-LNPs (+/- PEG-DMG) ***(Figure 7e and f)***. This reiterates the need for a safe and efficient transfection agent to maximize the uptake of siRNA in neurons, a hard-to-transfect cell type.

Around 94-96% of the cells were Cy5 (+) for the Lipofectamine RNAiMAX group indicating efficient uptake ***(Figure 7b right and f)***. Charge-based interactions of cationic lipids with negatively-charged cells allows for higher uptake as compared to neutral or negatively-charged particles. About 44-46% of the cells were Cy5 (+) for the naked Cy5 siRNA transfection group ***(Figure 7c (red))***. The difference between the uptake mediated by LNPs (+ PEG-DMG) and Lipofectamine RNAiMAX was statistically non-significant ***(Figure 7f)***. Despite the similar levels of uptake among the LNP(+PEG-DMG) and lipofectamine groups, it must also be pointed out that the uptake mediated by Lipofectamine RNAiMAX was accompanied with noticeable cell stress/toxicity upon visual observation under a microscope whereas LNPs showed a superior safety profile.

### Cy5 siRNA uptake by fluorescent microscopy in a human cortical neuron cell line

We studied the effect of exposure time on the uptake of LNPs by incubating the cells for two, four or 24 hours. PEGylation is known to regulate the uptake kinetics of LNPs into cells (49). We compared the differences in uptake for LNPs prepared with/without PEG-DMG. Although Lipofectamine RNAiMAX is a gold standard transfection agent for RNA molecules, it is also quite toxic to cells owing to its strong cationic nature. Therefore, it was unsurprising when we saw changes in morphology indicating cell death, just 2 h after Lipofectamine RNAiMAX was added to cortical neurons, while untreated cells continued to appear spindle-shaped and healthy ***(Figure 8a***).

For this experiment, cells were incubated either with naked Cy5 siRNA, Cy5 siRNA LNP with PEG-DMG, with Cy5 siRNA LNP without PEG-DMG, with Lipofectamine RNAiMAX Cy5 siRNA, or were left untreated. We qualitatively compared the fluorescence intensity among the cells treated with the above samples. Cells treated with naked Cy5 siRNA showed less intense fluorescent signals as compared to cells treated with Cy5 siRNA-LNPs (+/- PEG-DMG) for the two- and four-hour incubation time points ***(Figure 8a, b)***. However, we saw a noticeable increase in siRNA uptake 24 h-post transfection **(*Figure 8c*)**. The next pivotal observation was the difference in uptake from the LNPs (+/- PEG-DMG) groups. As mentioned earlier, PEG-DMG plays a key role in determining the physical stability of LNPs as it allows to maintain lower particle diameters by inhibiting particle aggregation (38). We also noted time-dependent differences in Cy5 siRNA uptake for the LNPs formulated with and without PEG-DMG. LNPs formulated with PEG-DMG showed a time-dependent increase in the uptake of Cy5 siRNA as opposed to LNPs without PEG-DMG where the extent of uptake was not time-dependent. We observed that only a few cells did not show Cy5 siRNA signals when treated with Cy5 siRNA-LNPs (+ PEG-DMG) two- and four-hours post-transfection whereas 24 h post-treatment almost all the cells showed signals corresponding to Cy5 siRNA. Conversely, cells treated with Cy5 siRNA-LNPs (- PEG- DMG) showed efficient uptake of Cy5 siRNA post two- and four hours transfection. A zoomed- in image of cells treated with LNPs containing 50 nM Cy5 siRNA is presented in ***Figure 8*d** to show image resolution. This particularly depicts the uptake of Cy5 siRNA LNPs associated with diffuse cytoplasmic fluorescence.

## Discussion

The goal of this pilot study was to determine if LNPs based on the benchmark C12-200 lipidoid can be used for siRNA transfection into neurons, a notoriously challenging yet clinically unmet/interesting delivery target. We propose that the well-documented delivery advantages of LNPs including their high levels of siRNA encapsulation, favorable pharmacokinetics, limited immune activation, uptake into target cells and efficient endosomal escape for siRNA into the cytoplasm (14, 47, 48) make them interesting candidates for neural delivery applications.

In this pilot study, we prepared and characterized C12-200 LNPs using DLS. The RiboGreen assay is a well-validated quantitative method to determine the % siRNA loading in LNPs (49, 50). Encapsulation of siRNA in the LNPs for the slow and fast mixing (without PEG-DMG) speeds were about 74.4% and 83.8%, respectively. LNPs with and without PEG-DMG (for fast mixing speed) showed encapsulation efficiencies of about 71.4 % and 83.8 %, respectively ***(Figure 1)*** which were comparable to the entrapment of siRNA reported for some of the top LNP candidates (54). Although we did not find a significant difference in the encapsulation efficiencies of LNPs prepared with and without PEG-DMG, the inclusion of PEG-DMG serves other vital roles *i.e.*, maintaining lower particle diameters by inhibiting particle aggregation and providing a stealth effect for longer circulation times *in vivo* (37, 38). We further studied the physicochemical stability of the prepared LNPs and compared the stabilizing effect provided by the inclusion of PEG-DMG and siRNA in the LNPs. We utilized siGFP as the model siRNA to compare the siRNA-loaded and blank LNP counterparts prepared both with and without PEG-DMG. We observed a non-significant increase or change in the particle diameters, polydispersity indices and zeta potential over a one-week storage period at 2-8 °C for all the samples **(Figure 2)**. Nevertheless, the particle diameters differed significantly for blank *vs*. siRNA-loaded LNPs and LNPs with *vs*. LNPs without PEG-DMG **(*Figure 2a*)**. A similar trend was also observed for the polydispersity indices of blank and siRNA-loaded LNPs; both with and without PEG-DMG **(Figure 2*b*)**. Our data demonstrated that both siRNA loading and PEG-DMG provided a stabilizing effect to the LNPs by maintaining lower particle diameters and uniform dispersity indices over a period of seven-days. The stabilizing effects of siRNA and PEG-DMG on the resulting particle diameters and dispersity indices were noted when the LNPs were stored for 24 h at 37°C **(*Figure 3*)**. LNPs showed further lower particle diameters (ca. 80.5 nm) when sizes were measured in the presence of serum (10% FBS) suggestive of the additional serum-mediated stabilization **(*Figure 4)***.

Although LNPs are deemed to be safe and are well-tolerated by most of the cells, determining their safety and tolerability in neural cells was our primary aim. As shown in ***Figure 6***, LNPs were deemed to be well-tolerated by primary DRG neurons as evident by >95% cell viabilities. Increasing the dose of siRNA (and concomitantly the dose of the lipids) did not alter the safety profile of LNPs ***(Figure 6c)***. Establishing the safety profile of LNPs with neural cells allowed further evaluation of their uptake in neural cell lines. We studied the uptake of Cy5 siRNA-loaded LNPs in primary rat cortical neurons and a human neuronal cell line using flow cytometry and fluorescence microscopy, respectively. Previously, Howard *et al* and Thiramanas *et al.* studied the cellular uptake of Cy5 siRNA using flow cytometry (52, 53). As PEG-DMG is known to stabilize the LNPs with resulting lower particle diameters and thus have an impact on their cellular uptake, we compared LNPs prepared with (+) and without (-) PEG-DMG. We used the siRNA complexes of the cationic transfection agent, Lipofectamine RNAiMAX, as a positive control. The cell-associated fluorescence (Cy5 (+) neurons) for the Cy5 siRNA-LNPs (+ PEG-DMG) group was about 94%—2-fold greater than the neurons transfected with naked Cy5 siRNA group **(Figure 7*c, e*, and *f*)**. As expected, despite its transfection-induced toxicity, Cy5 siRNA-Lipofectamine RNAiMAX-treated cells demonstrated the highest proportion of Cy5 (+) cells **(Figure 7*c, e, and f*)**. On the other hand, cells treated with LNPs appeared healthy and showed no visible signs of toxicity, highlighting the safety of LNPs over cationic transfection agents like Lipofectamine RNAiMAX. There was a significant difference in the uptake of neurons transfected with naked Cy5 siRNA as compared to the neurons transfected with the LNPs (both with and without PEG-DMG) emphasizing the need for an effective transfection agent for neural cell uptake **(Figure 7*f*)**.

We also performed fluorescence microscopy to qualitatively study the uptake of Cy5 siRNA into neural cells. A technical caveat of such qualitative assessments is rooted in the fact that the observed/apparent fluorescent intensities are not normalized to cell number. The cells appeared to be healthy, and spindle-shaped in all the treatment groups except for the Lipofectamine RNAiMAX group where cells appeared stressed and rounded as early as at the 2 h timepoint ***(Figure 8a)***. As described earlier, we speculate that this is due strong cationic nature of Lipofectamine RNAiMAX. We did not observe a greater uptake for cells treated with LNPs at 100 nM siRNA concentration **(*Supplementary Figure 2)***. Nearly all the cells in the field showed Cy5 fluorescent signals 24 h-post exposure **(*Figure 8c)*** compared to the 2 and 4 h time points **(*Figure 8a and b)***, suggesting that neural cells require a longer transfection time compared to non-neural cells. LNPs formulated with PEG-DMG showed a time-dependent increase in the uptake of Cy5 siRNA as opposed to LNPs without PEG-DMG. The time dependent uptake of LNPs formulated with PEG-DMG was in agreement with the PEG-DMG dissociation kinetics as predicted by Mui *et al*. (57). The PEG-lipid component of the LNPs undergoes desorption from the surface of LNPs which results in efficient uptake of the cargo into the cells (58). Studies have reported desorption rates of PEG-DMG from the LNPs at a rate of 45%/h only after which LNPs were taken up by cells (57). We observed a similar trend of Cy5 siRNA uptake when neurons were transfected with LNPs (+ PEG-DMG). At the 2 and 4 h time points, most (but not all) of the cells had efficiently taken up the LNPs. However, 24 h post-treatment, almost all the cells in the field of view showed fluorescent puncta corresponding to Cy5 siRNA. In contrast, in cells transfected with Cy5 siRNA-LNPs (- PEG-DMG), all the cells demonstrated complete and uniform uptake irrespective of the incubation period. The effects of RNAiMAX-mediated toxicity were evident via altered cell morphology seen in the phase contrast images.

Overall, both flow cytometry and fluorescent microscopy analysis in cortical neurons demonstrated efficient uptake of siRNA delivered using LNPs in the absence of noticeable toxicity while the cationic Lipofectamine RNAiMAX-mediated uptake was concomitant with significant cellular toxicity. Based on these findings, we conclude that LNPs are a safe carrier for siRNA delivery to neural cells. We are currently screening a pre-existing LNP library prepared using different lipidoid chemistries to identify LNP candidates for safe and efficient neuronal gene knockdown. We anticipate that the results of these studies will set the foundation for using LNPs for neural cell transfection in a variety of CNS diseases. While our approach validates using LNPs for uptake of siRNA into neural cells, their knockdown efficacy and therapeutic index remain an open question which we are currently investigating.

## Conclusion

SiRNA-loaded LNPs remained stable over a period of seven days and were well-tolerated by neural cell models, allowing their further exploration for gene silencing. Although the LNPs are comparable to Lipofectamine RNAiMAX in facilitating siRNA uptake into cortical neurons, the ionizable cationic LNPs are safer and well-tolerated compared to the cationic Lipofectamine RNAiMAX. PEG-DMG served a crucial role in maintaining lower particle diameters that likely resulted in efficient uptake into primary rat cortical neurons. Our findings suggest that LNPs hold promise for further silencing of therapeutically-relevant neuronal targets like TRPV1.

## Acknowledgements

This work was supported via start-up funds for the Manickam laboratory from Duquesne University (DU), a 2021 Faculty Development Fund (Office of Research, DU) and a 2021 Charles Henry Leach II Fund to the PI (Duquesne University). We are grateful to Dr. Kathryn A. Whitehead (Carnegie Mellon University) and her group for their support. We are grateful to Drs. Jane E. Hartung and Michael S. Gold (University of Pittsburgh) for providing the DRG cultures. We are also thankful to Dr. Wilson Meng and express our special appreciation to Mr. Nevil Abraham (Duquesne University) for his help with the qPCR experiments. We express our appreciation to Mr. Duncan Dobbins (Duquesne University) for his assistance with the graphical abstract.

## Response to reviewers

<u>We thank the reviewers for their insightful comments and helpful suggestions. We have addressed the comments and the individual responses to the questions are presented below.</u> All new text (and figure titles) added in the revised manuscript in response to the reviewer’s comments are highlighted in blue font.

**Reviewer #1: The authors give a specific neural application of LNPs which is a hot topic during the COVID-19 pandemic. There are some points to clarify before publication:**

**1. Although a reference is cited in the section “Preparation of siRNA-loaded LNPs”, this section can be described in more detail for the ease of understanding of readers out of the field. These parameters could be detailed: i) precise volume of ethanolic phase and aqueous phase used for the fabrication of LNPs should be cleared, ii) the ’slow mixing’ and for the ’fast mixing’ protocols should be explained in detail, iii) how the ethanol is removed from LNPs before application on the cells.**

The precise volumes of the aqueous and ethanolic phases have been added to the revised manuscript in **Table 1**. We have also appended the protocols to include details of the ‘slow’ and ‘fast’ mixing in the methods section under ‘**Preparation of siRNA-loaded LNPs (siRNA-LNPs)**’. We treated cells with non-dialyzed LNPs and the concentration of ethanol in the transfection/treatment mixture was ca. 2% v/v. In our personal correspondence with Dr. Kathryn Whitehead (Carnegie Mellon University) ca. 2018, one of the pioneers in the development of siRNA-loaded LNPs, she indicated that non-dialyzed LNPs could be safely used on cells as long as the cells tolerated the formulations with no significant toxicity. We observed ∼100% cell viabilities when cancer cell lines (MCF-7, MDA-MB-231 and BT-549) and primary DRG neurons were exposed to LNPs (**Figure 6**) and conclude that the non-dialyzed LNPs are safely tolerated by the cell models used in this study.

**2. The colloidal stability of nanoparticles should be also checked at 37 C for 24 h since the cells were exposed to nanoparticles for 24 hours in 37 C. Testing the stability at 37 C reveals if the nanoparticles remain stable during this experiment. Additionally, it would be nice to have the results of the serum stability test results of LNPs to see if the developed LNPs are stable in this medium.**

The colloidal stability of LNPs at 37°C for 24 h and in the presence of serum was studied and the results are presented in the revised manuscript as **Figures 3 and 4**. Our data showed that the LNP diameters and zeta potentials did no change appreciably 24 h post-preparation at 37 °C and LNP particle diameters were additionally stabilized in the presence of 10% FBS—suggesting that both these conditions did not affect their colloidal stability.

**3. TEM imaging before and after incubation at 37 C for 24 h could be presented to prove the stability of LNPs.**

As discussed above, we have used DLS measurements as a quantitative tool to demonstrate the stability of LNPs. We thank the reviewer for this important suggestion and are now establishing collaborative arrangements with the University of Pittsburgh to carry out cryo-TEM studies of LNPs and the results of these studies will be reported in a forthcoming manuscript.

**4. In figure4, some of the graphs and the letters on these graphs are presented with the same color which makes it harder to follow and understand. This figure could be rebuilt with more clear structuring and coloring. Specifically, the graph lines with green color seem pixelized.**

This figure has been re-worked to improve clarity in the revised manuscript (**Figure 7**). We would like to highlight a few things to explain the pixelization of the histogram. The green-colored histogram in **Figure 7** shows the percentage of positive cells for Cy5 (Cy5 (+)) when the neurons were treated with Cy5 siRNA/Lipofectamine RNAiMAX complexes. As expected, Lipofectamine RNAiMAX showed the highest (94.9 %) percentage of Cy5 (+) cells. As a result, neurons treated with Lipofectamine RNAiMAX showed the greatest shift towards the right. Interestingly, we also observed two distinct peaks (a “bimodal” histogram) for this group with varying Cy5 intensities as compared to a “unimodal” histogram for neurons treated with LNPs (**Figure 7c (yellow and brown)**). As RNAiMAX is toxic to cells, it is likely that the first peak corresponds to Cy5 siRNA taken up by distressed cells (resulting in lower Cy5 intensities). The second peak showed a greater percentage Cy5 (+) cells demonstrating the increased uptake of Cy5 siRNA by the viable cells. The histogram corresponding the Lipofectamine groups appears to be pixelized as the histogram width is also broader for this treatment group compared to the LNP-treated neurons. Moreover, the histograms depicted in **Figure 7** are autogenerated by the Attune NxT software of the Attune NxT Flow Cytometer. It should be pointed out that histograms reported in literature as well as those in the training and user guides of the Attune NxT Cytometer show a similar pattern of pixelization (1–4>).

**5. The fluorescent microscopy images presented in Figure5 seem blurry, they could be enhanced. Additionally, in Figure5b the cell morphology seems different than other cells specifically in the image of cells exposed to RNAİMAX complex (Cells treated with this complex are presented as viable after 4 h, it would be necessary to detail this finding).**

The ‘blurriness’ of the fluorescent microscopy image presented in **Figure 8** (in the revised manuscript) is likely a result of the ‘zoomed-out’ micrographs presented in the figures. We have revised **Figure 8** by including a ‘zoomed-in’ version of one the LNP groups (cells treated with 50 nM Cy5 siRNA-LNPs+PEG-DMG in **panel d**. This image depicts a clearer view of the cells wherein the Cy5 siRNA LNPs show diffuse fluorescence in the cytoplasm. The reviewer has rightly pointed out the difference in morphology of cells exposed to Lipofectamine RNAiMAX in panels b and c in **Figure 8**. Lipofectamine RNAiMAX is a highly cationic lipid and a benchmark transfection agent for RNA molecules. Although a strong cationic charge serves to enhance the uptake of carrier molecules in cells, it concurrently mediates cell stress (5). <u>The observed changes in morphology of cells is a result of this Lipofectamine-mediated stress in this particular cell model, human cortical neurons.</u>

**Reviewer #2: The author herein demonstrate performance of LNPs for siRNA delivery to neural cell. The study is interesting and deemed fit for publication in this journal. However, there are several limitations that must be addressed and discussed.**

**1. The author must perform a reporter assay. Only uptake does not guarantee successful siRNA delivery.**

The reviewer makes an important comment here and we have indeed performed pilot experiments to study the knockdown of a therapeutic/neuron-relevant knockdown target. We had chosen not to include this data in the original submission due to the reasons discussed in the supplementary information, but this data is now included in the revised manuscript (**Supplementary Figure 3**). Primary dorsal root ganglion (DRG) neurons isolated from rat trigeminal ganglia were used to study the knockdown of transient receptor potential cation channel subfamily V member 1 (TRPV1), a neuron-relevant knockdown target. SiRNA against TRPV1 (siTRPV1) was chosen as a proof-of-concept drug because it has been reported that TPRV1 becomes hyperactive in response to chronic inflammatory pain that reduces their threshold for activation and increases sodium, calcium and chloride fluxes (7, 8). TRPV1 is a non-selective cation channel exhibiting high calcium permeability and is expressed in peripheral, central axon terminals in the spinal cord, C fibers and/or Aδ fibers (8). Nearly 60% of the peptidergic primary nociceptors in the dorsal root ganglia and trigeminal ganglia express TRPV1 (8). Primary DRG cultures were isolated using previously reported methods (9). The inherent technical challenges associated with isolating DRG neurons/cultures resulted in low numbers of isolated cells but we still proceeded with the transfection study to determine if this TRPV1 target can be silenced using siTRPV1 delivered via LNPs (10–12).

**<u>A key limitation of this study is that the low numbers of neurons (2,500 cells/well) used will naturally have a low (or rather a very low) baseline expression of TRPV1 and therefore, this current setting does not allow us to optimally determine the effectiveness of siTRPV1 delivery via LNPs</u>**. Proceeding with this caveat, we cautiously discuss here the findings from this experiment. siTRPV1-LNPs (+PEG-DMG) were formulated using C12-200, an ionizable cationic lipid and helper lipids. The resulting mRNA levels post-transfection were determined using quantitative reverse transcription PCR. Our data showed a low 9% knockdown of TRPV1 when primary DRG cultures were treated with siTRPV1-LNPs that was similar to cells treated with the positive control, siTRPV1-Lipofectamine RNAiMAX complexes. Cells treated with naked siTRPV1-LNPs showed around 2.6% knockdown of TRPV1 whereas inverted (inv.) siTRPV1- LNPs showed about 3.2% TRPV1 knockdown. Despite the low levels of knockdown, the observed differences in % knockdown were significant (****p < 0.0001). As stated earlier in this section, this pilot study must be carefully interpreted due to the caveats associated with the low cell numbers and therefore, a lower baseline TRPV1 expression. Nevertheless, this data points out the safety and the potential of LNPs as delivery agents to silence therapeutically-relevant neuronal targets. Current efforts are underway in the laboratory to establish primary neuronal cultures with a higher yield and results of those studies will be reported in a forthcoming manuscript.

**2. The authors just selected one concentration to show cytocompatibility. The cytocompatibility of the LNPs and control lipids must be performed in a dose escalation setting for better comparison.**

The cytocompatibility data of LNPs tested at increasing doses of siGFP LNPs is now included in the revised manuscript (Figure 6c).

**3. The intensity and autocorrelation function of DLS measurement must be included. This reviewer strongly recommends comparing autocorrelation function at different time point to get insights into their stability.**

The intensity and autocorrelation function of DLS measurement is now included in the revised manuscript in Figure 4. We have also compared the autocorrelation function at different time points and is now included in the revised manuscript as Table 2.

**4. The author must stain the live cells. Besides, the microscopy pictures are of poor resolution.**

The ‘blurriness’ of the fluorescent microscopy image presented in Figure 8 (in the revised manuscript) is likely a result of the ‘zoomed-out’ micrographs presented in the figures. We have revised Figure 8 by including a ‘zoomed-in’ version of one the LNP groups (cells treated with 50 nM Cy5 siRNA-LNPs+PEG-DMG) in panel d. This image depicts a clearer view of the cells wherein the LNP-delivered siRNA shows diffuse fluorescence in the cytoplasm.

**5. The LNP preparation methods need clarification. How much volume of 1 mg/mL siRNA solution was prepared? The author must mention the cat# for each siRNA used.**

The precise volumes of the aqueous and ethanolic phases have been added to the revised manuscript in **Table 1**. We have also appended the protocols for ‘slow’ and ‘fast’ mixing in the methods section under ‘**Preparation of siRNA-loaded LNPs (siRNA-LNPs)**’. The catalog numbers of the siRNAs are now included in the ‘**Materials**’ section of the revised manuscript.

**6. In LNP preparation, why the author chose to dilute instead of dialysis/similar technique to replace the citrate buffer with PBS? What’s the final pH of the solution? This is important to report the zetapotential, as it is highly pH dependent. Depending on volume of citrate and PBS used for LNP preparation for different experiment their charge and size may get affected.**

In our personal correspondence with Dr. Kathryn Whitehead (Carnegie Mellon University) ca. 2018, one of the pioneers in the development of siRNA-loaded LNPs, she indicated that non-dialyzed LNPs could be safely used on cells as long as the cells tolerated the formulations with no significant toxicity. We observed ∼100% cell viabilities when cancer cell lines (MCF-7, MDA-MB-231 and BT-549) and primary DRG neurons were exposed to LNPs (**Figure 6**) and conclude that the non-dialyzed LNPs are safely tolerated by the cell models used in this study.

The final pH of the solution is 7.4 (now included in the legend for Figure 2 in the revised manuscript).

**7. The method section must include statistical analysis. Two-way ANOVA and not one-way should be performed whenever applicable.**

We have now added statistical analysis in the ‘**Methods**’ section of the revised manuscript. We used either one-way ANOVA or two-way ANOVA along with One-Sample T- and Wilcoxon tests based on the recommendations from GraphPad Prism software.

---

**Supplementary Figure S1.**
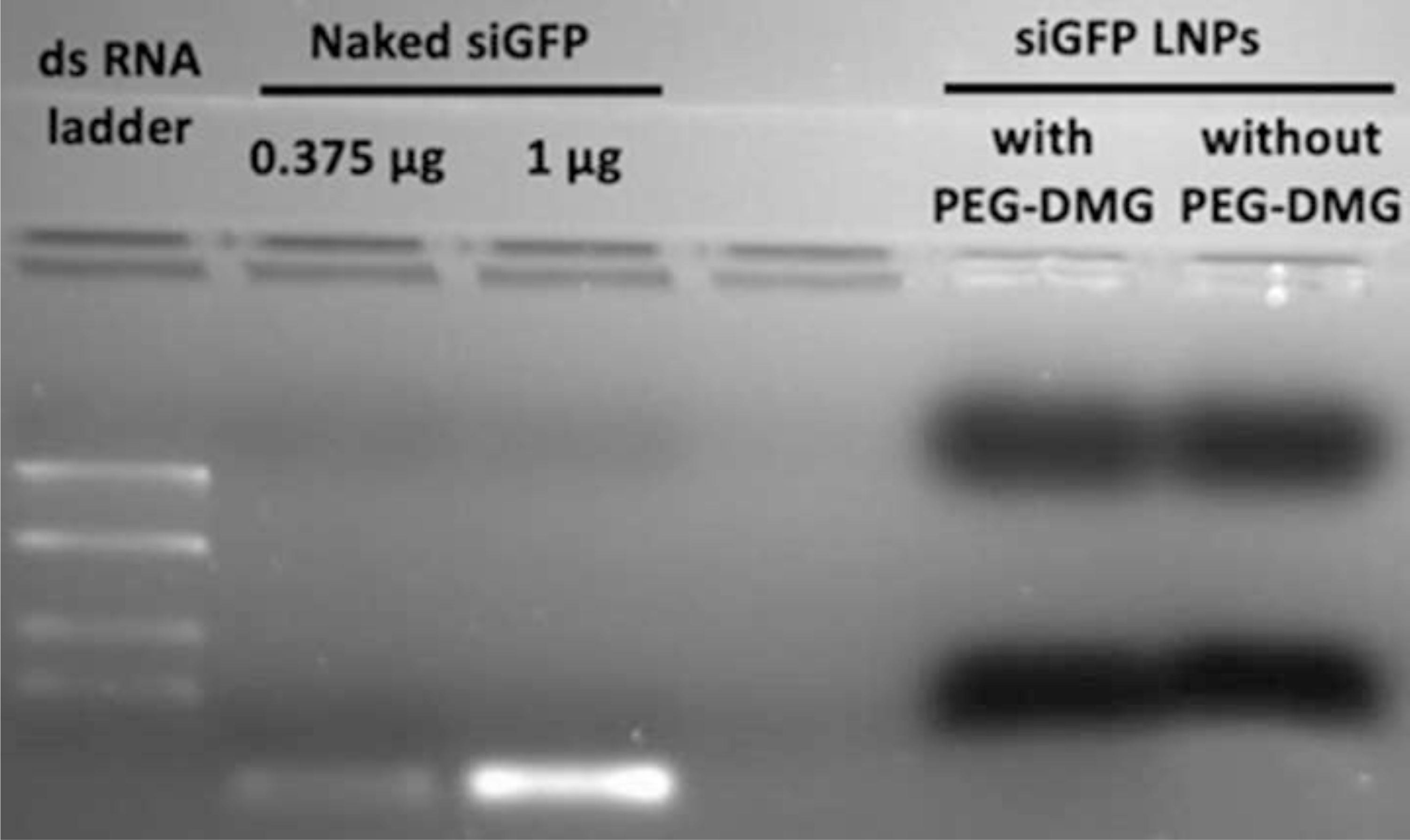

**Supplementary Figure S2.**
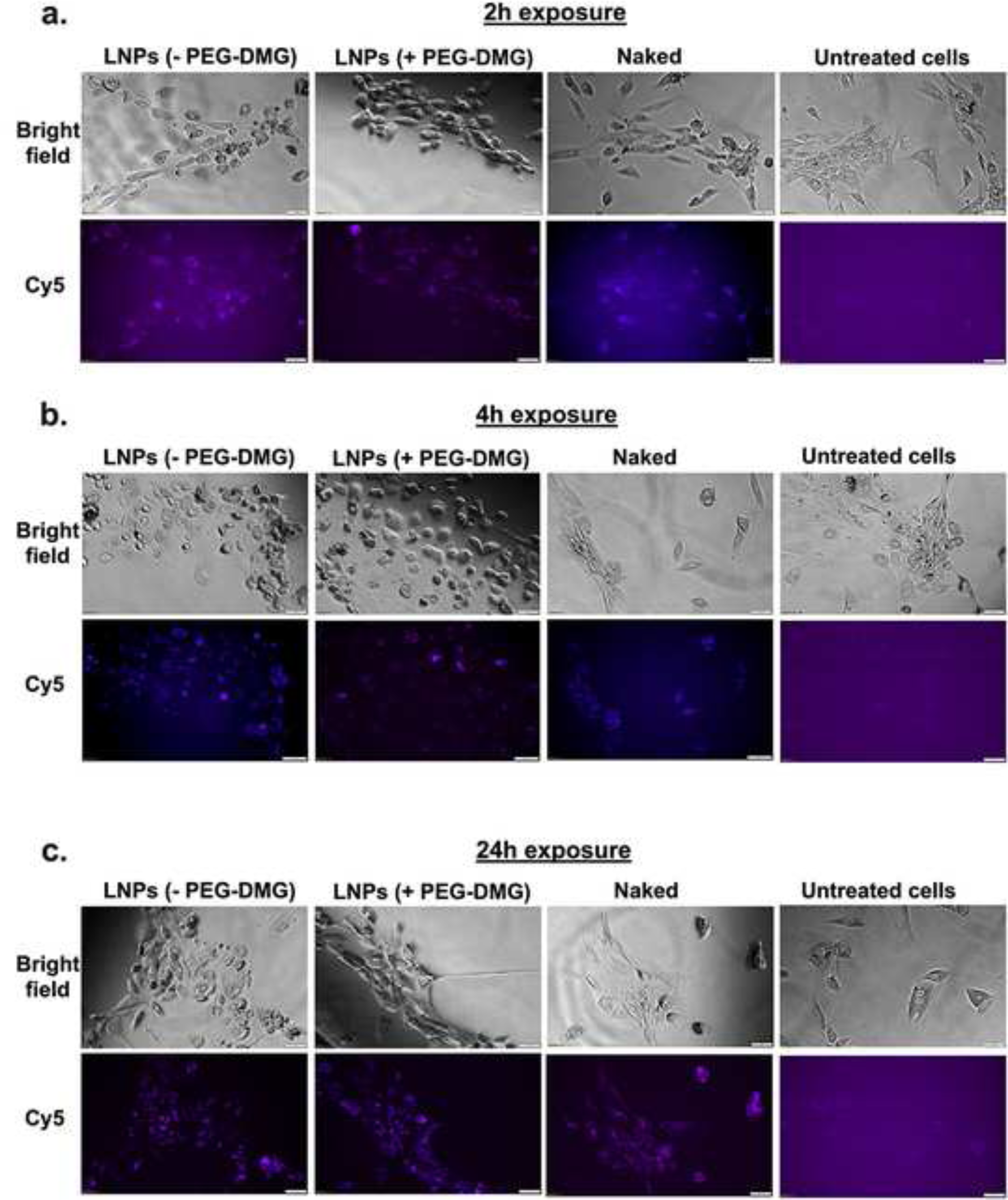

**Supplementary Figure S3.**
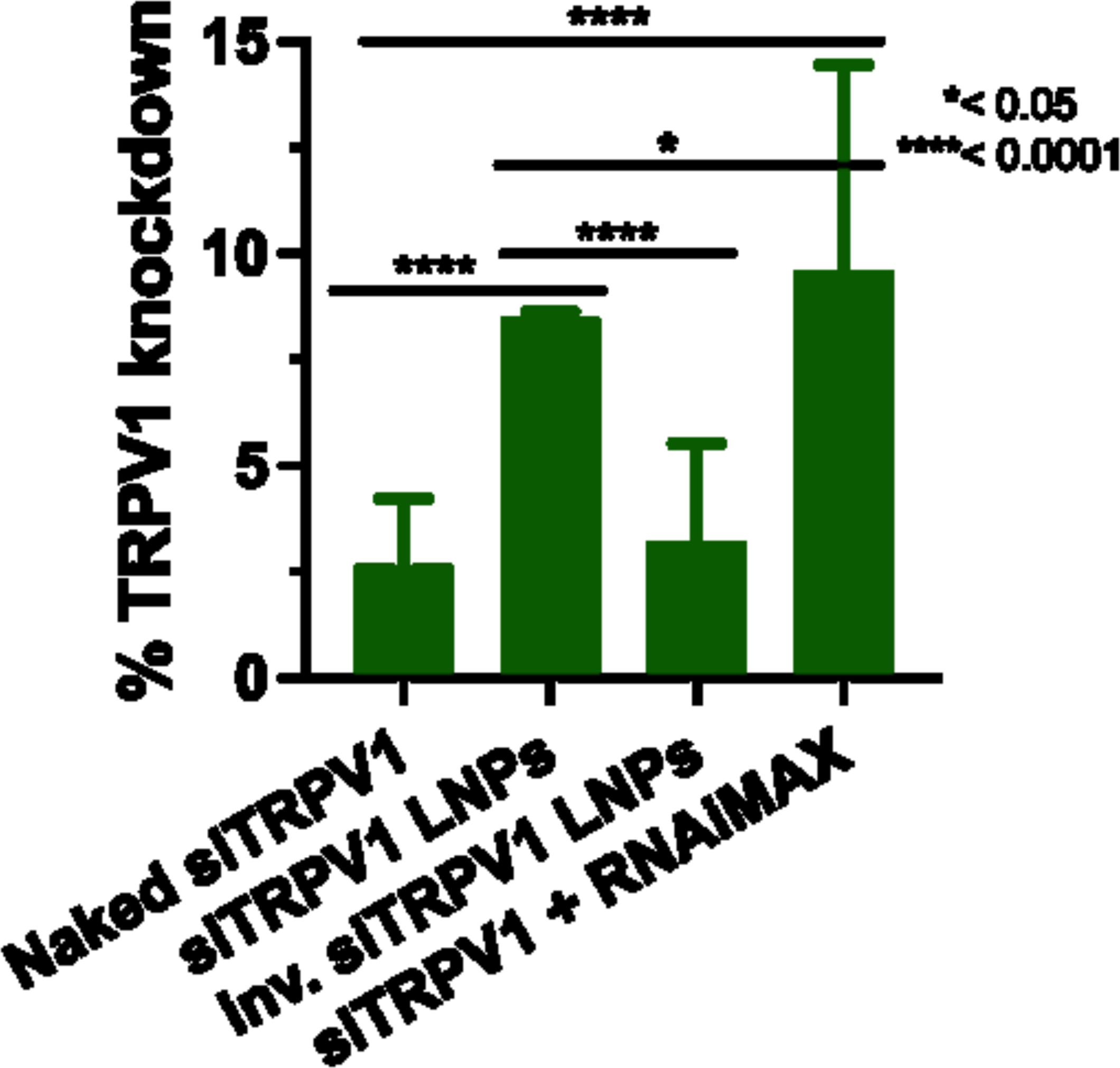

